# Modulation of pedunculopontine input to the basal ganglia relieves motor symptoms in Parkinsonian mice

**DOI:** 10.1101/2024.03.06.583786

**Authors:** Caixia Lin, Margreet Ridder, Jinhao Zhong, Eduardo A. Albornoz, Petra Sedlak, Li Xu, Trent M. Woodruff, Fangyi Chen, Pankaj Sah

## Abstract

Locomotion, the ability to move, is driven by central pattern generators in the spinal cord. However, its initiation, selection of gait and arrest are controlled by command centers in the brain stem of which the best understood is the mesencephalic locomotor region (MLR). Stimulation of the MLR can initiate or arrest movement, but its neural projections and the neural circuits that mediate these actions are poorly understood. Here we show that glutamatergic neurons in the pedunculopontine nucleus (PPN), a major part of the MLR, project to the substantial nigra pars reticulata (SNr) where they innervate neurons that send afferents to the ventro medial motor thalamus (VM). Stimulation of this input to the SNr in mice slows voluntary and driven movement by modifying gait, with higher frequencies arresting gait. In Parkinsons disease (PD), a neurodegenerative disorder, the cardinal symptoms are bradykinesia, tremor and rigidity. Using an mouse model of PD, we show that inhibition of PPN input to the SNr rescues bradykinesia. This is the first description of a circuit in the mammalian brain that regulates movement and gait selection. Targeting of this circuit may be a more effective modality for treating the motor symptoms of PD.

## Introduction

The ability to move is fundamental to the survival of all animals, and provides the means to explore the environment, catch prey and escape danger(*1*). Effective locomotion requires efficient transitions between the different phases of initiation, movement, and termination of gait (*2*). In quadrupeds, after intitiation of movement, coordinated patters of limb movement are engaged as needed. Thus, at lower, exploratory speeds, animals can either ‘walk’ or ‘trot’, whereas at higher speeds they ‘bound’ or ‘gallop’ where different gaits engage distinct sets of limb movement (*3, 4*). These patterns of movement are driven by specific neural circuits, the central pattern generators (CPGs) located in the spinal cord (*1, 2, 5*). However, the initiation, selection of gait, and termination of movement rely on supraspinal command centers (*1, 2*) and the basal ganglia (*1*).

Locomotor command centers are located in the brainstem, of which the best understood is the mesencephalic locomotor region (MLR) (*1*). First identified as a region in which electrical stimulation evoked locomotion in decerebrate cats(*6*), the MLR is present in all vertebrates(*1, 2, 5, 7*), and consists of the pedunculopontine (PPN) and cuneiform nucleus (CnF) (*2, 8*). Both nuclei are involved in locomotor control (*9–11*), but the neural circuits engaged are little understood. Thus, targeted optogenetic activation of glutamatergic neurons in the CnF can initiate high speed movement from rest, but the neural circuits which select the type of gait engaged are not known (*9–11*). The role of the PPN is less clear, and stimulation of neurons within the PPN have shown mixed results with some studies reporting increases in movement(*10*), some a reduction(*11, 12*), or arrest of movement(*13*) and others reporting no effect(*10*). Functionally, ascending PPN projections are thought to initiate goal directed movement via the motor cortex(*2, 9, 14*) while descending connections are thought to initiate motor arrest via the spinal CPGs(*13*). However, the neurons in the PPN that control these different behaviours, or their targets are not known.

The basal ganglia comprise a set of subcortical nuclei that play a central role in motor control, and disorders within the basal ganglia lead to a range of movement disorders such as Parkinsons disease (PD) (*15, 16*). The cardinal motor symptoms of PD, bradykinesia, tremor and rigidity have been long treated with deep brain stimulation within the subthalamic nucleus STN(*17*), also a component of the BG. However, in advanced PD, many patients also suffer from freezing of gait (FoG), in which individuals have a difficulty initiating gait(*18*) and deep brain stimulation of the PPN has emerged as a target to treat this disorder(*16*). How stimulation of the PPN relieves symptoms of PD is not clear. Here we show that a population of glutamatergic neurons in the PPN innervate neurons in the substantial nigra pars reticulata (SNr) that project to the ventromedial motor thalamus (VM). Stimulation of PPN input to the SNr reduces locomotor drive by modulating gait, and at high frequencies can arrest movement. Using a mouse model of PD that presents with depletion of dopaminergic neurons in the SNc (*19*), we show that the loss of locomotor drive in the model is rescued by inhibition of this PPN input to the SNr. These results define a specific neural circuit in the mouse brainstem that modulates locomotor drive by controlling gait. Our results provide an explanation for how stimulation in the PPN relieves the symptoms of PD and suggests more targeted interventions to control the symptoms of PD.

## Results

### PPN input to SNr is glutamatergic

The PPN is a heterogenous structure that contains local circuit GABAergic neurons and projection neurons that are either cholinergic or glutamatergic(*20, 21*). Tract tracing studies(*21*) have found reciprocal connections between the PPN and substantia nigra (SN), that is divided into the pars reticulata (SNr) and pars compacta (SNc). Electrophysiological recordings from the SNr neurons have found an excitatory input from the PPN(*22, 23*), and while both cholinergic and glutamatergic neurons send projections to the SNr (*24–27*), the identity of the excitatory transmitter is not known. To address this question we first injected retrograde adeno-associated virus (AAV) expressing tdTomato into the SNr (Fig. 1a, b), revealing extensive retrograde labelling of neurons in the PPN (Fig. 1c). Using GAD67-eGFP mice to label GABAergic neurons, and choline acetyltransferase (ChAT) immunohistochemistry to identify cholinergic neurons, we found no overlap between retrogradely labelled PPN neurons and cholinergic neurons (n = 3 mice; Fig. 1c-e), and only a very small number of retrogradely labelled neurons expressed eGFP (1.15 ± 0.99%; n = 3; Fig. 1c-e). By exclusion, PPN neurons that project to the SNr are glutamatergic. Retrogradely labelled cells were more abundant in the caudal PPN (n = 3, Fig. 1f), consistent with the distribution of glutamatergic neurons in the PPN(*20, 21*). To confirm the glutamatergic nature of the PPN input to the SNr, Cre-dependent AAV (EF1a-DIO-tdTomato) was injected into the PPN of Vglut2-Cre mice (n=3 mice) resulting in abundant labelling of glutamatergic neurons with tdTomato (Fig. S1a, b). tdTomato-positive axonal terminals originating from the PPN were found in the SN, with abundant labelling in the SNc and SNr (Fig. S1c, d), consistent with a glutamatergic projection from the PPN to the SNr. Notably, PPN glutamatergic neurons projected to both ipsilateral and contralateral SNr.

**Figure 1.**
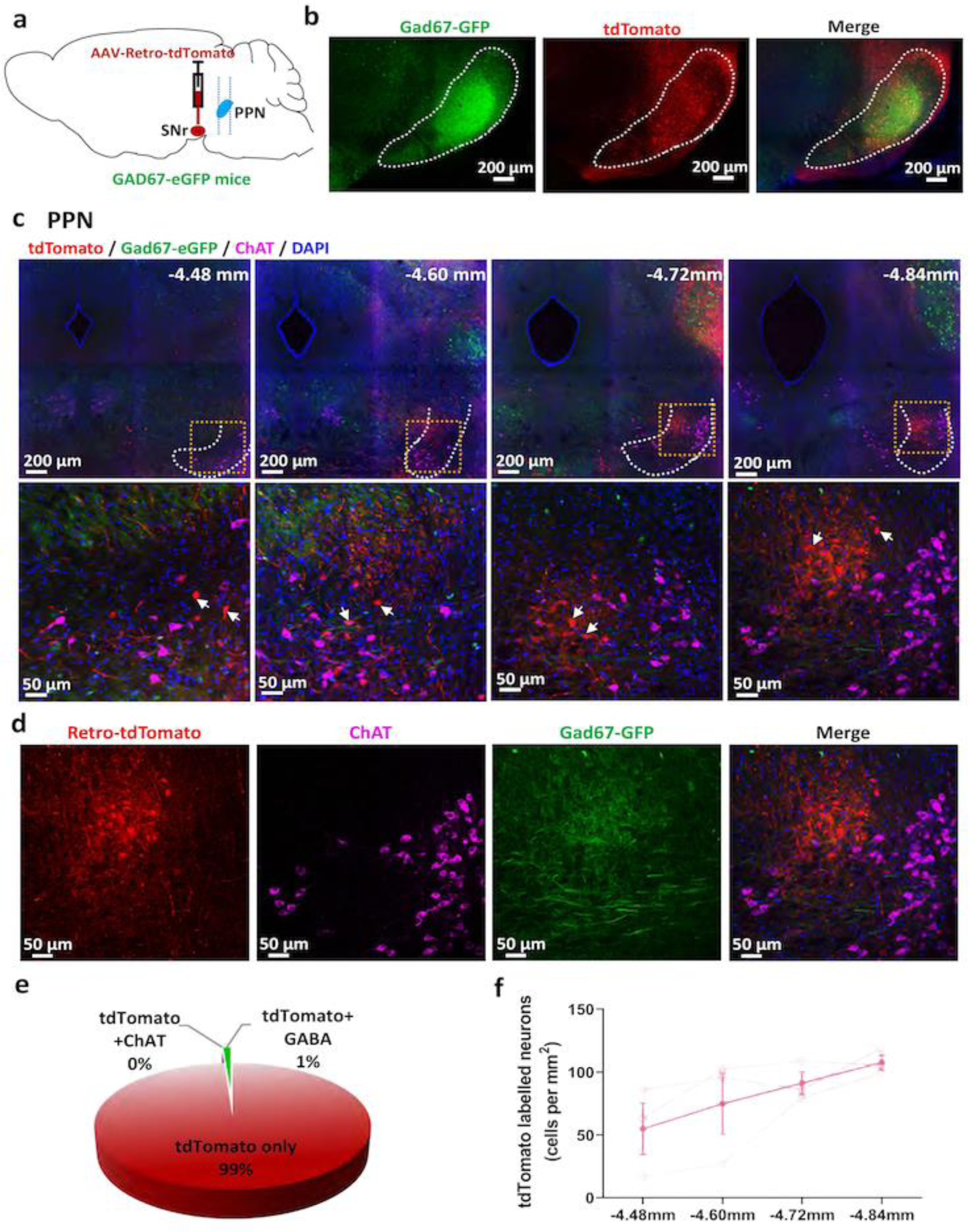
PPN input to the SNr is glutamatergic. **a,** Schematic illustrating procedure for backlabelling of PPN projection neurons to the SNr in GAD67-eGFP mice. **b,** Fluorescence images showing the injection site of tdTomato (red: tdTomato expressing cells) within the SNr (green: GAD67-eGFP positive cells; blue: DAPI). **c,** Top panels show the back-labelled retro-tdTomato positive neurons (red) in the PPN with sections from rostral to caudal immunolabelled for ChAT (purple) to mark cholinergic neurons. Locations of the brain sections are indicated with the light blue dashed lines in panel **a** (bregma −4.48mm to −4.84mm; PPN border depicted with white dashed line. bottom panels are enlargements of the yellow boxed areas of top panels, showing high resolution images of retro-tdTomato positive neurons (red, white arrowheads), ChAT neurons (purple) and GABAergic neurons (green, GAD67-eGFP) in the PPN; blue, DAPI. **d,** Representative fluorescence images showing the distribution of tdTomato expressing neurons (red), cholinergic neurons (purple, ChAT) and GABAergic neurons (green, eGFP) in the PPN with a merged image at Bregma −4.84 mm (blue, DAPI). **e,** Percentage of tdTomato single labelled, tdTomato double labelled with ChAT and tdTomato double labelled with GAD67-eGFP neurons in the PPN (quantified from 3 mice, n = 3). **f,** Distribution of tdTomato labelled neurons from rostral to caudal PPN. The percentage of tdTomato labelled neurons along the rostral-caudal axis was relative to total tdTomato labelled neurons in the whole PPN region (quantified from 3 mice, n = 3, error bars represent SEM).

To investigate the physiology of PPN input to the SNr, channelrhodopsin (ChR2) was selectively expressed in PPN neurons projecting to the SNr. Retrograde AAV expressing Cre-recombinase was injected into the SNr of wild-type C57BL/6J mice, and Cre-dependent AAV-DIO-ChR2-mCherry into the PPN (Fig. 2a). Four weeks following viral injection, abundant ChR2-mCherry positive neurons were seen in the PPN (Fig. 2b), and ChR2-mCherry labelled axon terminals from the PPN were present throughout the SNr (Fig. 2c). Using *ex vivo* brain slices prepared from injected animals, whole-cell recordings were obtained from neurons in the SNr (Fig. 2c). Blue light (470 nm) stimulation evoked excitatory synaptic currents (EPSCs) in 81% of SNr neurons (44 out of 54, Fig. 2d, 13 mice). PPN input to individual SNr neurons ranged from <10pA (Vh −60 mV) to almost 300pA (Fig. S2a, n=44), which drove action potentials in 60% of neurons (15 out of 25, Fig. 2e, 9 mice). These EPSCs had a tight response latency (3.54 ± 0.88 ms; n = 44; Fig S2b), and application of tetrodotoxin (TTX) blocked this input, and addition of 4-aminopyridine (4AP) rescued the photo-stimulation evoked response (Fig. 2f, n = 10) indicating that they were monosynaptic. Paired pulse stimulation (Fig. S2c, n = 4) revealed that the PPN input has a high release probability and a depressing nature. The evoked EPSC was abolished by the AMPA receptor antagonist 6-cyano-7-nitroquinovaline-2,3-dione (CNQX; 10 µM), confirming that PPN input to the SNr was glutamatergic (Fig. 2g, n = 8). Application of the nicotinic cholinergic receptor antagonist (mecamylamine; 10μM), had no effect (Fig. S2d, n = 4, *P* = 0.9166), and depolarization of SNr neurons did not evoke any outward synaptic current (Fig. S2e, n = 3), showing there is no fast cholinergic or GABAergic component to the PPN input. As a final confirmation we expressed ChR2 exclusively in glutamatergic neurons by injection of Cre-dependent AAV-DIO-ChR2-YFP into the PPN of Vglut2-Cre mice (Fig. 2h-j). Whole cell recordings from SNr neurons in *ex vivo* slices showed that most recorded SNr neurons (85%) received an excitatory input from the PPN that was blocked by CNQX (Fig. 2k, l).

**Figure 2.**
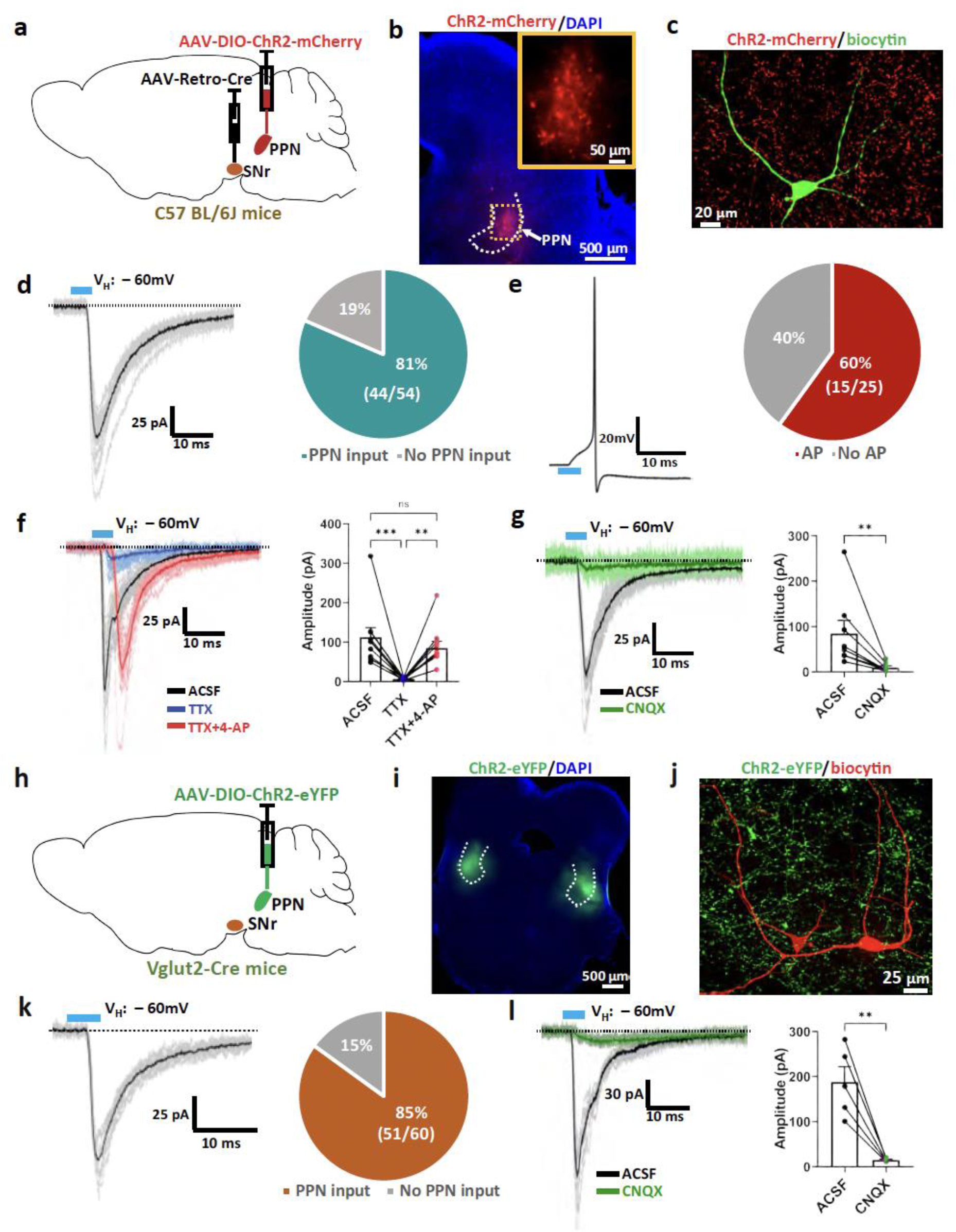
SNr neurons receive monosynaptic glutamatergic input from the PPN. **a,** Schematic illustrating labelling PPN to SNr specific projection neurons with ChR2. AAV-retro-Cre was injected into the SNr and AAV-DIO-ChR2-mCherry into the PPN of C57BL/6J mice. **b,** ChR2 labelled neurons (red) in the PPN 6 weeks post viral injection. **c,** A representative recorded neuron (green, biocytin filled) in the SNr surrounded by ChR2 terminals (red) originating from the PPN. **d,** Voltage-clamp recording from a representative SNr neuron shows the response to a 5 ms 470-nm light pulse (blue bar) at a holding potential of −60 mV (bold black: averaged response; light grey: raw traces; dashed black line indicates the baseline). Right pie chart displaying the percentage of SNr neurons (13 mice) responding to optical stimulation. **e,** Representative current-clamp recording of a single light stimulation response from an SNr neuron evokes an action potential. Right pie chart displaying the percentage of SNr neurons firing in response to optogenetic stimulation. **f,** Representative traces (left) and boxplots (right) showing PPN to SNr input is monosynaptic (n = 10). The light evoked EPSC (black, averaged response; light grey: raw traces) recorded at −60mV was blocked by tetrodotoxin (TTX, bold blue: averaged response; light blue: raw traces) and rescued by 4-AP (red: averaged response; light red: raw traces). **g,** Representative traces (left) at −60 mV from a representative SNr neuron in a C57BL/6J mouse were blocked by the application of CNQX (green: averaged response; light green: raw traces; dashed black line indicates the baseline), showing that the PPN input to SNr neurons is glutamatergic. Right bar graph displays the summary of the response amplitudes before and after CNQX application (n = 8). **h**, Schematic illustrating transduction of ChR2 in glutamatergic neurons in the PPN using AAV-DIO-ChR2-eYFP injected in the PPN of Vglut2-Cre mice. **i**, ChR2 labelled neurons (green) in the Vglut2-Cre mouse brain slices 6 weeks post viral injection (left panel). PPN border depicted with white dashed line. Blue: DAPI. **j,** Representative recorded SNr neurons (red, biocytin filled) from a Vglu2-Cre mouse surrounded by ChR2 terminals (green) originating from the PPN. **k,** Voltage-clamp recording of the light stimulation response at −60 mV from a representative SNr neuron in a Vglut2-Cre mouse (bold black: averaged response; grey: raw traces; dashed black line indicates the baseline). Right pie chart displaying the percentage of SNr neurons (12 mice) which respond to optogenetic stimulation in Vglut2-Cre mice. **l,** Light evoked EPSCs from a representative SNr neuron in a Vglut2-Cre mouse before (bold blue: averaged response; light blue: raw traces) and aftersCNQX application (red: averaged response; light red: raw traces; dashed black line indicates the baseline). Right plots show the summary of the response amplitudes before and after CNQX application (n = 5). For all panels, error bars represent SEM. ** p < 0.01, *** p < 0.005. “ns” indicates no significant difference.

### PPN input to SNr reduces locomotor speed by changing gait

Our electrophysiology and tract tracing results show that the excitatory PPN input to the SNr is glutamatergic, and we next asked what role it has in locomotor control. We began by testing the impact of increasing the activity of glutamatergic neurons in the PPN using Designer Receptors Exclusively Activated by Designer Drugs (DREADDs). Using Vglut2-Cre mice, the clozapine-N-oxide (CNO) activated excitatory DREADD, hM3DGq, was transduced in glutamatergic PPN neurons with AAV-DIO-hM3DGq-mcherry (Fig. S3a). Consistent with previous studies(*9–11*), we found that non-selective activation of glutamatergic PPN neurons induced locomotor promotion with mice receiving CNO spending more time moving (Fig S3e), and moving faster (Fig S3d) and further (Fig. S3b, c) in the open field. Mice treated with CNO also show a significantly higher latency to fall on the rotarod (Fig. S3f). Glutamatergic neurons in the PPN have a wide projection pattern with ascending afferents to the BG and descending input to the spinal cord(*2, 9*). To delineate the impact of ascending and descending projections we selectively stimulated excitatory into to the SNr. Retro-AAV-Cre was injected bilaterally into the SNr, and Cre-dependent AAV encoding hM3DGq (AAV-DIO-hM3DGq-mcherry) was delivered bilaterally into the PPN of C57BL/6J mice. This resulted in selective expression of hM3DGq in PPN glutamatergic neurons that project to the SNr (Fig. 3a). Mice were tested using two motor tasks: the open field (Fig. 3b) and rotarod. When PPN input to the SNr was selectively driven with CNO (see Methods) mice moved at slower speeds (Fig. 3c), generally spent less time moving (Fig. 3d), hence travelled for shorter distances (Fig. 3b e). There was no significant difference in the latency to fall off the rotarod (Fig. 3f). In contrast, when the activity of these neurons was reduced using the inhibitory DREADD, hM4DGi (Fig. S3g), mice injected with CNO travelled for longer distances (Fig. S3h, i) with a higher average locomotor speed (Fig. S3j) and spent more time moving (Fig. S3k). Similar to results using the excitatory DREADD hM3DGq, there was no change in the latency to fall in the rotarod test (Fig. S3l). Together these results indicate that stimulation of PPN input to the SNr slows locomotor speed.

**Figure 3.**
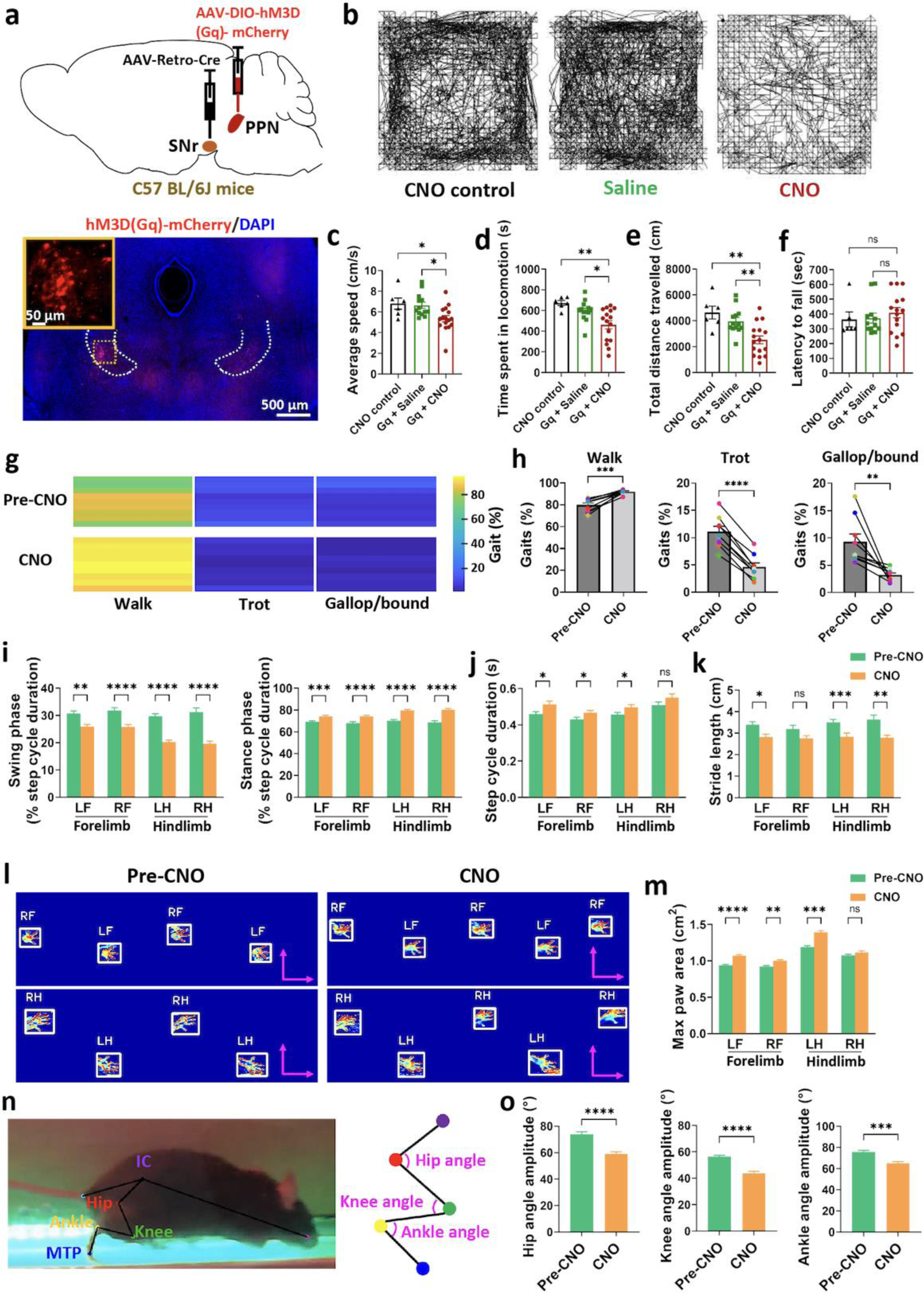
PPN input to SNr modulates locomotor activity by changing gait. **a,** Schematic illustrating retrograde labelling of PPN to SNr projection neurons with hM3D DREADDs using AAV-Retro-Cre and AAV-DIO-hM3D (Gq)-mCherry in C57 BL/6J mice. Bottom panel shows hM3D (Gq) DREADDs expressing neurons (red) in the PPN. PPN border depicted with a white dashed line; the insert shows the high-resolution image of hM3D (Gq) DREADD expressing neurons (red) in the PPN (enlargements of the yellow boxed area). Blue: DAPI. **b,** Representative moving path of mice from the CNO control group (left), saline group (middle) and CNO group (right) in the open field. **c - e,** Bar graphs show quantification of the distance travelled (**c**), velocity (**d**) and time spent in locomotion (**e**) in the open field of mice in the CNO control group (black, n = 6), saline group (green, n = 13) and CNO group (red, n = 16). Error bars represent SEM. **f,** Bar graph showing the latency to fall in the rotarod test of mice in three groups. Error bars represent SD. **g**, Locomotor gait patterns before (pre) and following (post) CNO injection with individual mice in separate horizontal lines (n = 9). **h**, Bar graphs showing quantification of different gaits (walk, trot and gallop/bound) before and after CNO application (n = 9). **i, j**, CNO application modified temporal gait dynamics, including swing phase and stance phase (**i**) and step cycle duration (**j**). **k,** Stride length was decreased by CNO. **l**, Footprints from a representative mouse before and after CNO injection. Scale bar, 1 cm. **m,** CNO application increased the maximum paw area of mice touching the treadmill surface. **n**, Hindlimb kinematics analysis by machine learning. Machine learning enabled markerless tracking of joint points (IC, hip, knee, ankle and MTP) on the mouse hindlimb. **o,** Mice treated with the CNO showed smaller angular variations of hip, knee and ankle joints on the hindlimb. For all panels except indicated, error bars represent SEM. * p < 0.05, ** p < 0.01, *** p < 0.005. **** p < 0.001. “ns” indicates no significant difference.

Kinematic analysis of locomotion in quadrupeds has led to the recognition of four different gaits that are associated with different speeds of movement, and patterns of limb movement(*3*). At slower speeds animals either ‘walk’ or ‘trot’, whereas higher speeds they can ‘gallop’ or ‘bound’ (*3*). We have shown that stimulation of PPN glutamatergic input to the SNr led to an overall reduction of movement while its inhibition led to an increase in movement. Kinematic analysis (see Methods) showed that stimuation of PPN input induced more ‘walk’ gaits while decreasing the higher speed ‘ trot’, ‘gallop’ and ‘bound’ (Fig. 3g, h), indicating that activation of the PPN to SNr pathway leads to a shift in locomotor gait. The reduction in movement is accomplished by a decrease in the step cycle duration (Fig. 3i) with a shorter swing and longer stance phase (Fig. 3j), that result in a shorter stride length (Fig. 3k, l) and smaller maximum paw area contacting the treadmill surface (Fig. 3m; Fig. S4). These changes in gait dynamics induced by CNO are consistent with the changes in the locomotor gait; moving from walking to gallop/bound gaits, is done by increasing the step frequency, and swing phase while decreasing the stride length (*3*). No significant changes in locomotor gait patterns (Fig. S5a, b) and gait dynamics (Fig. S5c-f) were seen in virally injected mice treated with saline.

To assess the influence of the PPN to SNr pathway on limb coordination, we investigated the changes in hindlimb kinematics using machine learning based markerless tracking of hindlimb movements from the side view while mice ran on a treadmill. Using markerless tracking, we positioned each hindlimb joint (iliac crest [IC], hip, knee, ankle, and metatarsophalangeal [MTP]) (Fig. 3n) and measured angular variations of the various joints. This analysis shows that the angular variations of the hip, knee and ankle joints decreased signicantly after CNO application, indicating that activation of the PPN-SNr input also modified limb kinematics, leading to hindlimb joint flexion(Fig. 3o).

### PPN input to the SNr can stop driven and voluntary locomotion

Simulation of neurons in the CnF has been found to stimulate speed to movement in a frequency dependent manner(*10, 11*). To test the effect stimulation frequency channelrhodopsin (ChR2) was transduced in glutamatergic neurons of the PPN by AAV-DIO-ChR2-YFP injection into the PPN of Vglut2-Cre mice. Optical fibers were bilaterally implanted over the SNr to stimulate glutamatergic PPN axon terminals in the SNr (Fig. 4a, Fig. S6a, b), and stimulation delivered over a range of frequencies to test the impact stimulating PPN input to the SNr while mice ran on a treadmill. Stimulation of PPN afferents to the SNr at 10Hz had little effect on movement (Fig. 4b), but increasing the stimulation frequency to 20Hz and 40Hz caused a reliable reduction of locomotion (Supplementary Video 1 and Video 2, available on request), with a complete halt of movement at 40 Hz (Fig. 4c, d). The mice displayed various locomotor gaits including ‘walk’ and ‘trot’ during the pre-laser period while slowing down or halting their locomotion immediately upon laser stimulation. Locomotor speed during laser stimulated was dramatically slower (Fig. 4c, d; Fig. S6c). The inhibitory effects were stronger with increasing laser power (Fig. S6d). There were no changes in locomotion speed and gait during the no laser trials, or in control mice (Fig. S6e, f). In the open field, when mice were freely moving, photo activation at 20Hz and 40Hz also led to the reduction of locomotion with a sharp reduction in the distance travelled and locomotor speed (Fig. 4e, f; Supplementary Video 3 and 4, available on request). Furthermore, in both tests, increasing the frequency of stimulation increased the inhibitory effect on locomotion. Laser stimulation lead to a stop in most trials at 40Hz (Fig. 4d, f) but only in half of the trials at 20Hz (Fig. 4c, e). Together with the chemogenetic manipulation, our data show that glutamatergic input from the PPN to SNr can reduce locomotion and stop movement by changing gait patterns.

**Figure 4.**
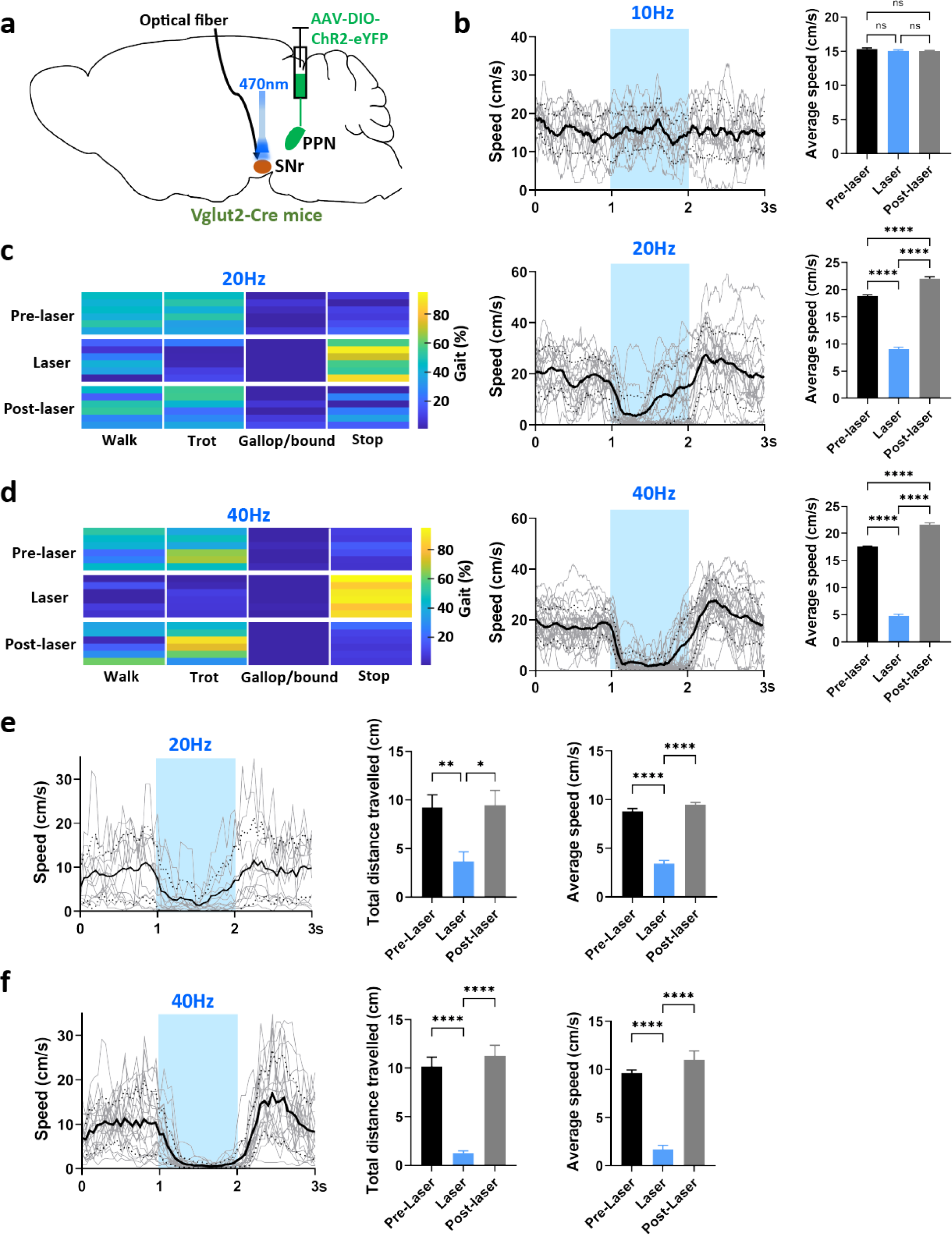
Frequency dependent modulation of gait by PPN afferents to the SNr. **a,** Schematic illustrating experimental configuration. AAV-DIO-ChR2-YFP was bilaterally injected into the PPN in Vglut2-Cre mice and optical fibers implanted bilaterally above the SNr. **b-d**, The effect of laser stimulation at 10 Hz (**b**), 20 Hz (**c**) and 40 Hz (**d**) while mice walked on a treadmill (Methods). **b,** Photo stimulation at 10 Hz has little effect on treadmill locomotion. Left panel shows the real-time speed versus time with single trials (in gray, 6 mice) and their average (in black). Blue region depicts the laser stimulation period. Right panel shows the average speed during three episodes: Pre-laser, Laser and Post laser (n = 6). **c,** 20 Hz stimulation significanlty slowed movement of mice running on the treadmill. Left panel shows the locomotor gait patterns during three episodes (Pre-laser, Laser and Post laser) with individual mice in separate horizontal lines (n = 6 mice). Middle panel shows the real-time speed versus time with single trials (in gray, 6 mice) and their average (in black). Blue region depicts the laser stimulation period. Right panel shows the average speed during 3 episodes: Pre-laser, Laser and Post laser (n = 6). **d,** 40 Hz photo stimulation halted movement of mice on the treadmill. Left panel shows the locomotor gait patterns during three episodes (Pre-laser, Laser and Post laser) with individual mice in separate horizontal lines (n = 6 mice). Middle panel shows the real-time speed versus time with single trials (in gray, 6 mice) and their average (in black). Blue region depicts the laser stimulation period. Right panel shows the average speed during 3 episodes: Pre-laser, Laser and Post laser (n = 6). **e, f,** The effect of laser stimulation at 20 Hz (**e**) and 40 Hz (**f**) on freely moving mice in the open field test. **e,** 20 Hz stimulation significantly slowed locomotion of the freely moving mice. Panels from left to right show the real-time speed (single trials in grey and average in black), total travelled distance and average speed during 3 episodes: Pre-laser, Laser and Post laser (n = 6). **f,** 40 Hz laser stimulation stopped freely moving mice. Panels from left to right show the real-time speed (single trials in grey and average in black), total travelled distance and average speed during 3 episodes: Pre-laser, Laser and Post laser (n = 6). For all panels, error bars represent SEM. * p < 0.05, ** p < 0.01, **** p < 0.001. “ns” indicates no significant difference.

### PPN to SNr input modulates locomotion via the ventromedial thalamus

The SNr is the major output of the BG motor network, with projections to multiple downstream regions(*28*). We have shown that stimulation of glutamatergic input from the PPN to the SNr acts as a brake to movement. To understand the downstream projections that mediate this effect we used a transsynaptic viral tracing strategy. The anterograde transsynaptic AAVs expressing Cre-recombinase (AAV1-hSyn-Cre) and AAV-DIO-YFP were injected into the PPN of C57BL/6J mice. Simultaneously, Cre-dependent AAV-DIO-tdTomato was delivered into the SNr (Fig. 5a, b). This approach labels neurons in the PPN with YFP, and as AAV1 spreads trans-synaptically in an anterograde direction(*29*), it enables the Cre-recombinase to spread to postsynaptic neurons in the SNr and drive expression of tdTomato in neurons which receive PPN input (Fig. 5b). By tracking the tdTomato positive terminals, the downstream targets of the PPN-SNr projection were uncovered. Afferent projections from PPN targeted SNr neurons were distributed to a number of key motor regions (Fig. 5c-e; Fig. S7a-c), including the ventromedial part (VM) of the motor thalamus, several BG nuclei, the subthalamic nucleus (STN) and the internal globus pallidus entopeduncular nucleus, (GPi), and other brain areas involved in motor control including the parafascicular nucleus (Pf) which has been recently reported to promote movement (*30*). These projections are known targets of the SNr, with the VM being a major target in the motor network (*28*). Our findings suggest that neurons in the SNr that project to the VM receive glutamatergic input from the PPN. To confirm this result, using Vglut2-Cre mice, retrogradely transported fluorescent beads were injected into the VM and AAV-DIO-ChR2-eYFP delivered into the PPN to transduce ChR2 in glutamatergic neurons (Fig. 5f). Following recovery, *ex vivo* brain slices were prepared and whole cell recordings obtained from bead positive neurons in the SNr that project to the VM. Stimulation of the PPN input confirmed that most (84.8%) VM projecting neurons in the SNr receive excitatory input from the PPN (Fig. 5g-j). Thus, glutamatergic excitatory input from the PPN innervates neurons in the SNr that project to the VM.

**Figure 5.**
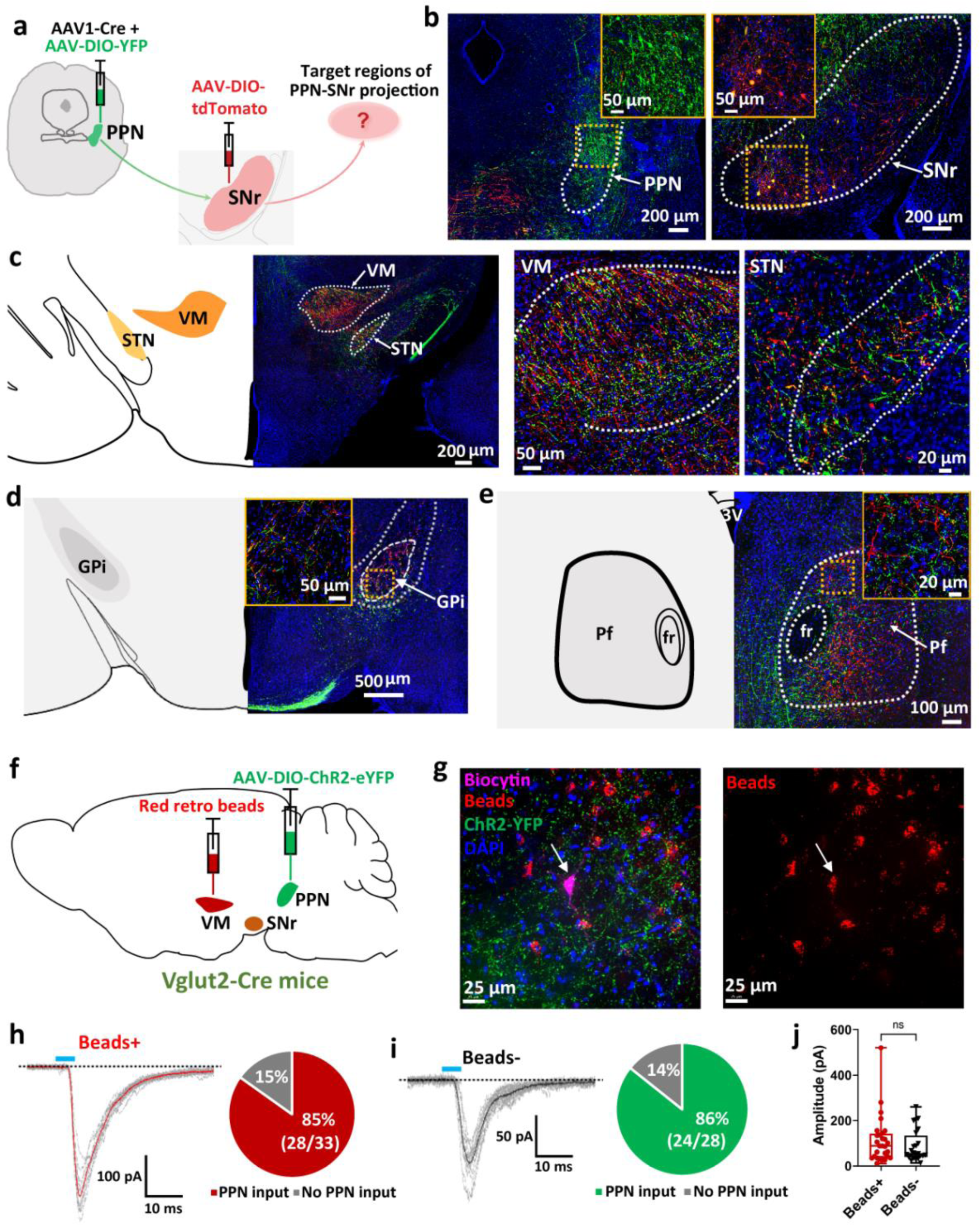
SNr neurons that receive PPN input project to the motor ventro medial thalamus and basal ganglia. **a,** Schematic illustrating procedure for antegrade transsynaptic tracing study. Injection of AAV1-Cre + AAV-DIO-YFP into the PPN allowed transsynaptic spread of Cre from presynaptic PPN into postsynaptic SNr. AAV-DIO-tdTomato was subsequently injected into the SNr resulting in the visualization of SNr neurons receiving PPN input. **b**, Fluorescence images showing the injection sites. Left panel shows a representative image of the PPN (green: YFP expressing cells, blue: DAPI). Right panel shows a representative image of the SNr (red: tdTomato expressing cells, blue: DAPI). Border of the PPN and SNr depicted with white dashed line. The yellow boxed inserts show enlargements of the yellow dashed boxed areas. **c-e,** Coronal sections showing terminal labeling sites of PPN-defined SNr neurons. **c,** Terminals of PPN-defined SNr neurons distributed extensively in the VM and STN. Border of the VM and STN depicted with white dashed lines; Right two panels show high magnification of the VM and STN separately. VM: ventral medial thalamus; STN: subthalamic nucleus: **d,e,** Terminal distribution of projections of SNr neurons receiving PPN input in the GPi and the Pf. GPi and Pf border depicted with white dashed line. The yellow boxed inserts show enlargements of the yellow dashed boxed areas. GPi: internal globus pallidus entopeduncular nucleus; Pf: parafascicular nucleus. **f,** Schematic for labelling glutamatergic neurons in the PPN by injecting AAV-DIO-ChR2-eYFP into the PPN and labelling VM projecting SNr neurons by injecting red retrograde beads into the VM of Vglut2-Cre mice. **g**, Reconstruction of a representative patched beads+ SNr neuron (green: ChR2; blue: DAPI; red: beads; purple: biocytin filled SNr neuron). **h,** Voltage-clamp recording of the light stimulation responses at −60 mV from a Beads+ SNr neuron (red: averaged response; grey: raw traces; dashed black line indicates the baseline). Right pie chart showing the percentage of beads positive neurons (7 mice) that respond to optogenetic stimulation. **i,** Voltage-clamp recording of the light stimulation response at −60 mV from a beads-negative SNr neuron (black: averaged response; grey: raw traces; dashed black line indicates the baseline). Right pie chart displaying the percentage of Beads-neurons (7 mice), which respond to optogenetic stimulation. **j,** Boxplots of the amplitude of the responses from Beads+ (n = 28) and Beads- (n = 24) neurons. Error bars represent SD.

The VM plays a key role in motor control(*14, 31*), and we therefore tested if locomotor control that the PPN exerts via the SNr also requires the VM. Using Vglut2-Cre mice, ChR2 was expressed in glutamatergic neurons in the PPN. Simultaneously, retrogradely transported AAV transducing Cre-recombinase was injected into the VM, and AAV-DIO-hM4D(Gi) was delivered into the SNr to transduce the inhibitory DREADD hM4DGi specifically in the VM projecting SNr neurons (Fig. 6a, Fig. S8b). Optical fibers were implanted bilaterally over the SNr to drive PPN input to the SNr (Fig. S8a). Following a period of recovery, animals were tested on the treadmill and open field. We tested the impact of stimulating PPN input to the SNr, before and after silencing SNr input to the VM using CNO (Fig 6). Electrophysiological recordings from the hM4DGi expressing SNr neurons *ex vivo* showed that CNO application decreased the firing rate and suppressed the AP response to light stimulation of the PPN ChR2 terminals (Fig. S8d,e), confirming the effect of the CNO. As shown above, photo stimulation of PPN input to the SNr (20Hz and 40Hz) led to a reliable reduction of locomotion (Fig. 6b-e). When SNr neurons projecting to the VM were silenced, the effect of PPN stimulation was significantly reduced (Fig 6b-e; Supplementary Video 5 - 8, available on request). Indeed, the reduction in locomotion in response to 20 Hz stimulation was completely blocked in the presence of CNO (Fig 6b, d; Supplementary Video 5 and Video 7, available on request), and was fully restored when CNO was washed out (Fig. S8c). These experiments show that the VM is necessary for the PPN to SNr projection suppressing locomotion, verifying that the PPN to SNr pathway is involved in the locomotor control neural circuits through the VM.

**Figure 6.**
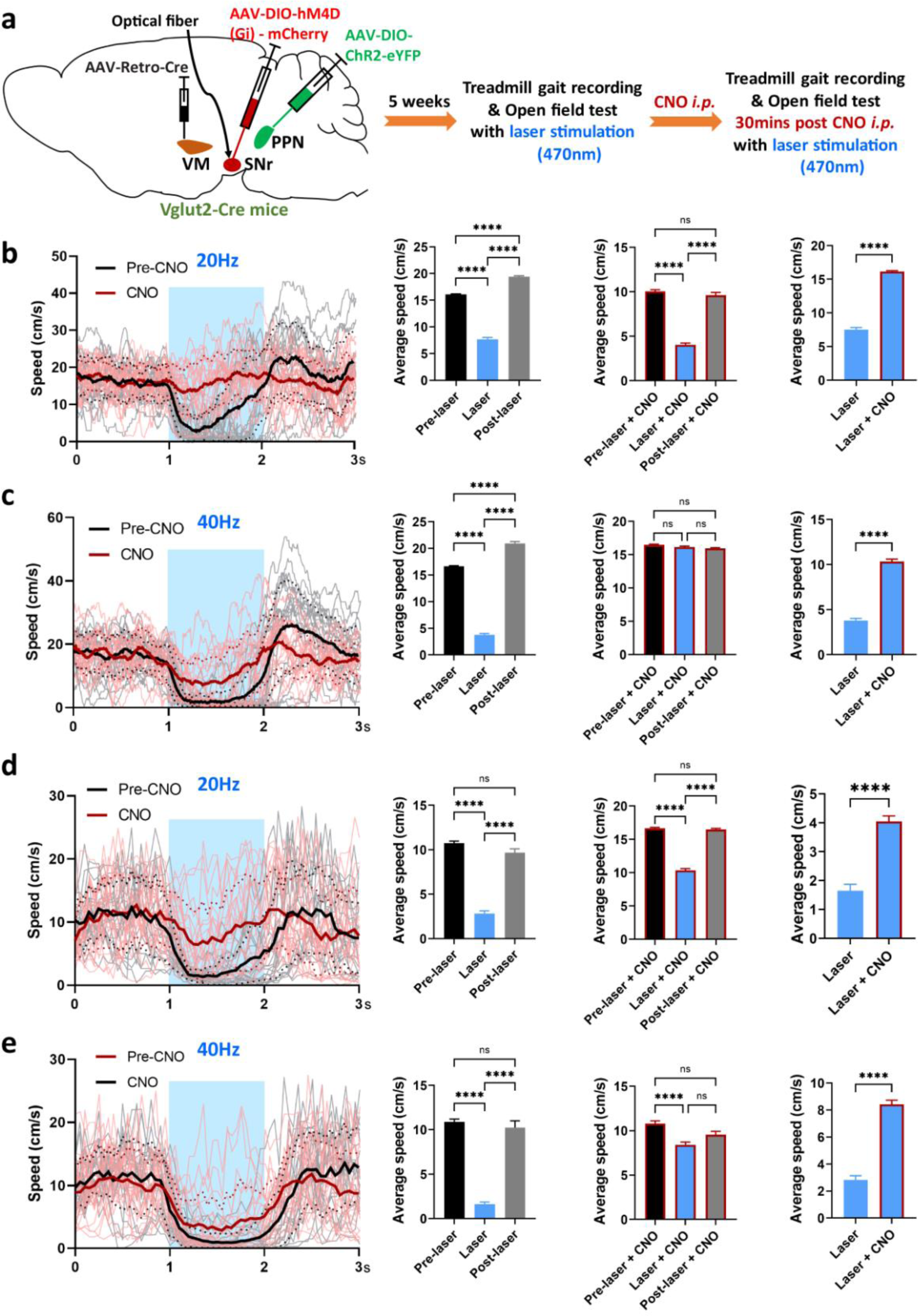
Input from PPN to the SNr inhibits locomotion via the ventro medial thalamus (VM). **a,** Schematic illustrating the procedure of behavioral experiments combined with optogenetics and chemogenetics. AAV-DIO-ChR2-YFP was bilaterally injected into the PPN and dual optical fibers were implanted bilaterally above the SNr in Vglut2-Cre mice. The inhibitory DREADD hM4D (Gi) was selectively expressed in the SNr to VM projection neurons with AAV-Retro-Cre injection into the VM and injection of AAV-DIO-hM4D (Gi)-mCherry into the SNr. 5 weeks post-surgery, behavioral experiments were performed with 470nm laser stimulation before and after CNO application. **b, c,** Effect of laser stimulation on mice running on the treadmill before and after CNO injection (n = 5). Panels from left to right - real-time speed (Pre CNO application: single trials in grey and average in black; post-CNO application: single trials in light red and average in bold red), average speed during 3 episodes (Pre-laser, Laser and Post laser) before and after CNO application, and average speed during laser pre and post CNO injection (n = 5). **b,** CNO application abolished the inhibitory effect on locomotion caused by 20 Hz laser stimulation. **c,** CNO application weakened the inhibitory effect on locomotion caused by 40 Hz laser stimulation. **d, e,** Effects of laser stimulation on freely moving mice in the open field test before and after CNO injection. CNO application weakened the inhibitory effect on voluntary movement of mice induced by 20 Hz (**d**) and 40 Hz (**e**) laser stimulation (n = 5). Panels from left to right - real-time speed (Pre CNO application: single trials in grey and average in black; post-CNO application: single trials in light red and average in bold red), average speed during 3 episodes (Pre-laser, Laser and Post laser) before and after CNO application, and average speed during laser pre and post CNO injection (n = 5). For all panels, error bars represent SEM. * p < 0.05, ** p < 0.01, **** p < 0.001. “ns” indicates no significant difference.

### Inhibition of PPN input to the SNr rescues motor deficits in PD mice

In PD, apart from loss of dopaminergic neurons in the SN, these is also significant loss of cholinergic neurons in the PPN(*32*), and low frequency stimulation of the PPN has emerged as a target for the amelioration of the motor symptoms of PD(*16, 33*). As activation of the PPN input to the SNr suppresses movement and can arrest locomotor gait, we hypothesized that inhibition of this pathway may have therapeutic effects on parkinsonian motor dysfunction. An animal model of Parkinsons was generated by unilateral injection of 6-hydroxydopamine (6-OHDA) into the striatum (CPu) (*34*) (Fig.7a), which led to the loss of dopaminergic neurons in the SNc (Fig. 7b) and diminished terminals in the CPu (Fig. S9a). To silence the PPN-SNr projection, archaerhodopsin (ArchT) was selectively expressed in PPN neurons projecting SNr neurons by bilateral injections of AAV-Retro-pAM-Cre in the SNr combined with Cre-dependent AAV-DIO-ArchT-GFP into the PPN of wild-type C57BL/6J mice (Fig. 7a, Fig. S9a). Dural optical fibers were implanted over the SNr to optogenetically inhibit the PPN input to the SNr (Fig. S9b). After two weeks’ recovery, mice tested on the rotarod had a significant decrease in the latency to fall off the rod in 6-OHDA injected animals compared with saline in injected mice (Fig. S9c). We next performed open field testing and treadmill recordings with laser stimulation. Consistent with the chemogenetic data (Fig. S3g-l), optogenetic inhibition of PPN input to the SNr increased motor activity and locomotor speed of saline-treated control mice in the open field (Fig. 7c). As expected, 6-OHDA mice displayed PD-like symptoms, including a shorter distance travelled and a reduced locomotor speed in the open field test. Inhibition of PPN input to the SNr led to a dramatic rescue of the motor deficit with a significant increase in the distance travelled and locomotor speed (Fig. 7d; Supplementary Video 9, available on request). Notably, the distance travelled by the PD mice during laser stimulation was similar to that of the saline control mice at baseline (Fig. S9d), showing that inhibition of the PPN - SNr projection reversed the loss of motor activity in PD mice. In contrast to saline control mice, the 6-OHDA mice exhibited some gait impairments including decreased swing phase, increased stance phase and shorter stride length, which were recovered by optogenetic silencing of PPN neuron terminals in the SNr (Fig. 7e). Overall, our data indicates that optogenetic manipulation of the PPN input to the SNr is sufficient to rescue the motor deficits in PD mice.

**Figure 7.**
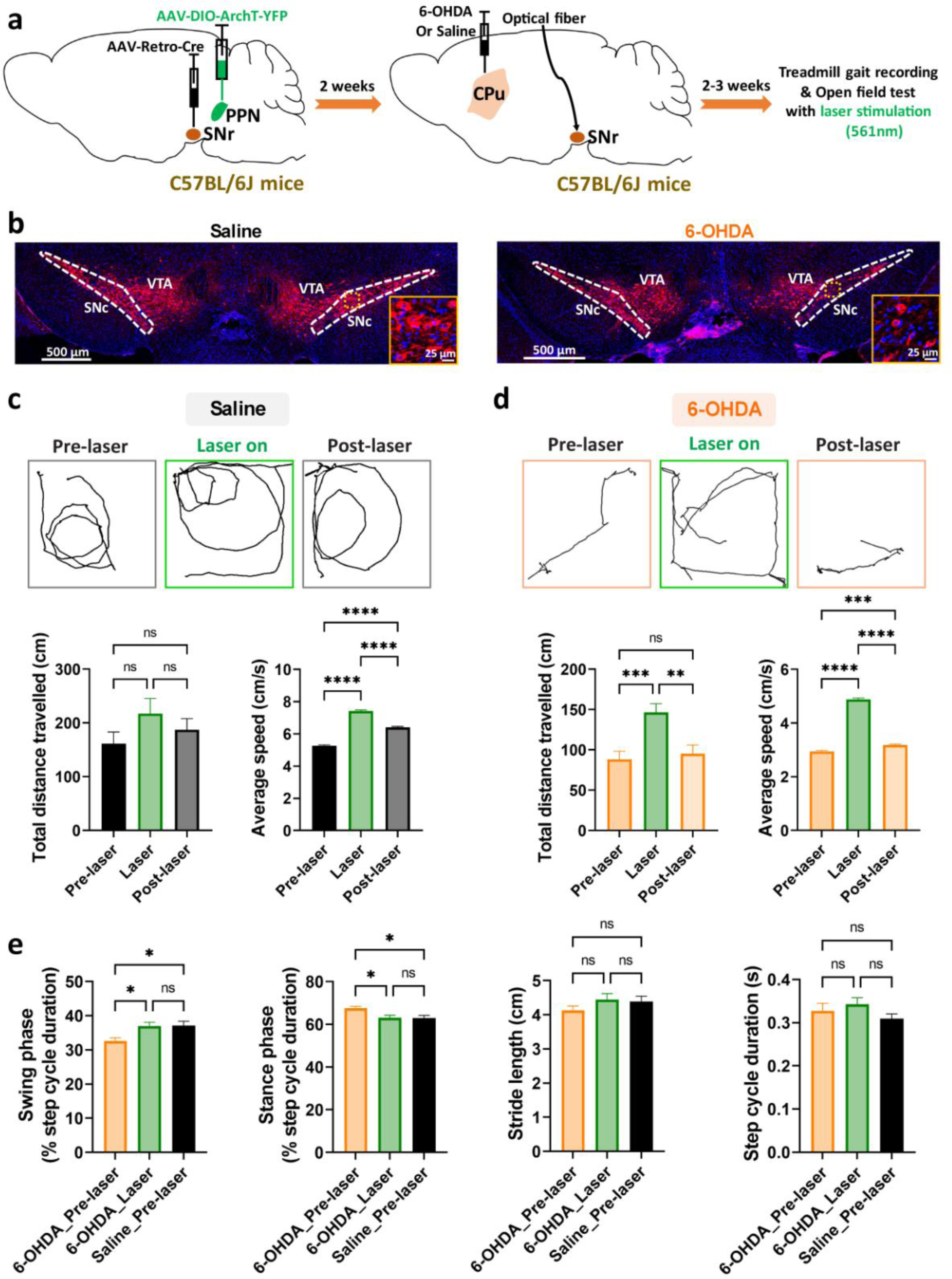
Optogenetic inhibition of the PPN input to the SNr rescues motor deficits in PD mice. **a,** Schematic of the surgery procedures and behavioural experiments. AAV-Retro-pAM-Cre was injected into the SNr and AAV-DIO-ArchT-YFP into the PPN of C57BL/6J mice. After 2 weeks later, 6-OHDA or saline was injected in right striatum (CPu) and dual optical fibers were implanted bilaterally above the SNr. 2-3 weeks post-surgery, behavioral tests were performed with 561nm laser stimulation. **b,** Representative images of TH labeled (red) dopaminergic neurons in the SNc of saline and 6-OHDA treated animals, showing loss of dopaminergic cells in the SNc after 6-OHDA injection in the right hemisphere. Blue: DAPI. **c, d**, Laser stimulation (561nm, 30s) increased movements of both saline-treated (**c,** n = 4) and 6-OHDA-treated mice (**d,** n = 6) freely moving in the open field test. Top panels show moving path of a representative animal during 3 episodes: Pre-laser, Laser and Post laser. Bottom panels show quantification of the total distance travelled and average speed during the three episodes. **e,** Laser stimulation modified gait dynamics of 6-OHDA treated mice running on the treadmill (6-OHDA treated mice, n = 4; saline treated mice, n = 3). Panels from left to right show comparation of gait parameters including swing phase, stance phase, stride length and step cycle. For all panels, error bars represent SEM. * p < 0.05, ** p < 0.01, *** p < 0.005, **** p < 0.001. “ns” indicates no significant difference.

Previous studies reported that following dopamine depletion in PD animal models, SNr neurons displayed abnormal firing activities along with changes in synaptic inputs to this region (*35–37*). To investigate whether the PPN input to the SNr was modified in PD mice, AAV-DIO-ChR2-EYFP was injected into the PPN combined with AAV-Retro-pAM-Cre into the SNr of wild-type C57BL/6J mice (Fig. S10a). 6-OHDA was injected unilaterally in the striatum for generating the PD mouse model (Fig. S10a, b). Two weeks post 6-OHDA injection, firing properties and light responses (470nm, 5s) were recorded from the SNr neurons by patch-clamp recordings. SNr neurons in 6-OHDA treated PD mice and saline control mice displayed similar firing frequencies (Fig. S10c). There were no significant differences in the amplitudes and PPRs of the light responses recorded from SNr neurons in 6-OHDA treated mice and saline treated mice (Fig. S10d, e) although the response latency was decreased in 6-OHDA treated mice (Fig. S10f), suggesting that the physiology of the PPN-SNr projection might not be modified in the PD mouse model.

## Discussion

The ability to move and explore the environment and escape danger are essential for survival (*1, 2, 5*). Active movement engages a repertoire of stereotypical movements driven by CPGs in the spinal cord, but the initiation, selection of gait, and its termination are controlled by circuits in the midbrain and BG(*2, 4, 5*). The MLR, consisting of the PPN and CnF is a key region involved in motor control(*1, 5, 38*). Within the MLR, activation of glutamatergic neurons in the CnF can initiate fast movement from rest, but the role of the PPN is less clear(*9–11*). Moreover, how activity in the MLR controls movement, and the neural circuits engaged, is not well understood. Despite this, the PPN has emerged as a target for the treatment of PD, a neurodegenerative disorder where bradykinesia is a cardinal symptom(*16*).

Neurons in the PPN and CnF have extensive distributions that include rostral and caudal motor areas(*10, 11, 27*). The SN is a key part of the midbrain motor control circuit and dysfunction within the SN leads to disorders like PD(*16*). Tracing studies have shown that the PPN projects to both the SNc and SNr(*21*), we have shown that the projection to the SNr is entirely glutamatergic. This result is in accordance with a recent study showing a subpopulation of MLR glutamatergic neurons projecting to the SNr(*27*). The SNr is the main output nucleus of the BG network, sends inhibitory output to upstream and downstream motor areas(*28*), with a major target being the VM motor thalamus(*28, 39*). Our results show that SNr neurons that receive PPN input, project to the VM. Stimulation of this input to the SNr, combined with machine learning based multi-animal pose tracking provided us with an efficient and accurate way to analyze the locomotor gaits and limb activity in both treadmill running and freely moving mice. This analysis shows that PPN input to the SNr input modulates locomotor speed and limb kinematics with an overall reduction in locomotion, an effect that is mediated via the VM.

While there is general agreement that the PPN is strongly involved in locomotor control(*1, 2, 5*), studies on its impact on locomotion have reached different conclusions(*10–12, 24*). In large part this may result from the fact that while previous studies have targeted neurons within the PPN, there has not been a separation of neurons with distinct targets. Our results identify a population of glutamatergic neurons in the PPN that project to the SNr, control the locomotion by effective modulation of gait. A recent study has described a population of glutamatergic neurons in the PPN defined by expression of the transcription factor Chx10(*13*). High frequency stimulation (40 Hz) of these neurons in the PPN resulted in motor arrest as well reduction of respiration and heart rate(*13*) and suggested that these cells define a circuit for global motor arrest(*13*). Low frequency stimulation was not tested, and the pathways that mediate changes in gait, respiration and heart rate were not examined. We suggest that the effects on gait evoked by stimulation of Chx10-positive PPN neurons results from a population that project to the SNr. Interestingly, the VM also receives direct glutamatergic projections from the PPN(*14*). Whether population of glutamatergic neurons in the PPN that project to the VM and those that provide disynaptic input via the SNr is an interesting question for future studies

Deep brain stimulation of the PPN has emerged as a target for the treatment for freezing of gait but the mechanisms of action remain unknown. Stimulation of PPN input to the SNr reduces movement and at high frequency halts locomotion, and both actions are mediated via projections to the VM. Bradykinesia is a key symptom of PD, and late stage PD patients can suffer from freezing of gait. Here, we show that inhibition of PPN input to the SNr rescues the bradykinesia in a mouse model of PD.

Overall, our findings demonstrate the existence of a glutamatergic projection from the PPN to the SNr and unravel a novel motor pathway connecting the midbrain, BG network and motor thalamus. Furthermore, we show that activation of distinct groups of PPN neurons may yield diverse, even opposite, effects on movement. The PPN has been used as a target for deep brain stimulation (DBS) for the treatment of PD(*16*), however, the efficacy of this therapy is under some debate with divergent results(*40, 41*). Our results suggest that if DBS targets distinct groups of PPN neurons, some may improve PD motor symptoms while some may have no effect or even exacerbate freezing in PD patients. Thus, further studies are still required to uncover the complicated mechanisms by which the PPN and related neural circuits modulate movement, in order to improve the clinical efficacy of PPN-DBS in PD patients. Together, this study reveals a novel neural circuit of the PPN for locomotor control, but also suggests the importance of developing more selective DBS strategies, which may lead to more reliable clinical outcomes for PD therapy.

## Methods and materials

### Animals

Male and female C57BL/6J, Vglut2-IRES-Cre (Jax 016963), and GAD67-eGFP mice were housed in a 12/12 h light/dark cycle with ad libitum access to food and water. All Vglut2-IRES-Cre mice (denoted Vglut2-Cre) used in this study were homozygous and GAD67-eGFP mice were heterozygous. Vglut2-Cre mice express Cre recombinase driven by vesicular-glutamate transporter 2 (Vglut2) promoter that is restricted to glutamatergic neuron cell bodies. GAD67-eGFP mice express enhanced green fluorescent protein (eGFP) under the control of the mouse Gad1 (Glutamate decarboxylase 1, GAD67) gene promoter. All experimental procedures were performed in agreement with the guidelines of the University of Queensland Animal Ethics Committee.

### Stereotaxic surgery

Mice (4-5 weeks) were anesthetized with 10% ketamine and 5% xylazine (10 ml/kg, i.p.) and placed in a stereotaxic frame. Incisions were made on the midline of animal’s head to expose the skull and to identify bregma and lambda. Small burr holes were drilled and virus or beads were injected using borosilicate glass pipettes (GC150F, 1.5 mm, Harvard Apparatus, UK) pulled with a PC-10 electrode puller (Narishige, Japan). When injecting, the pipette was connected to a picospritzer (Picospritzer II, Parker, US) using plastic tubing and air pressure was applied for 5-10 ms. The coordinates (all in mm related to bregma) used for PPN stereotactic injections in the mice were (anterior/posterior, medial/lateral, dorsal/ventral): −4.6, ±1.2, −3.5. Coordinates for the SNr injections were (A/P, M/L, D/V): 3.3, ±1.5, −4.5. Coordinates for the VM injections were (A/P, M/L, D/V): 1.4, ±0.8, −4.2. For each injection, the glass pipette was kept in place for 10min to limit the diffusion of the virus/beads. The incision was stitched with sterile sutures and closed using the tissue adhesive (Vetbond) at the end of the surgery. Metacam (0.4 μl/g) diluted in 1 ml of saline was injected subcutaneously. During all surgical procedures, mice were placed on a heating pad (37°C) and put back in their cages until they were fully conscious and were autonomous.

For anterograde tracing studies, adult Vglut2-Cre mice (P28-35) were injected with 100 nl AAV1/2-EF1α-DIO-tdTomato (3.49e12 GC/ml, produced in-house) unilaterally into the PPN. For retrograde tracing, adult GAD67-eGFP mice (4-6 weeks) were injected into the SNr unilaterally with 100 nl AAV-Retro-pAM-tdTomato (3.96×10e11 GC/ml, produced in-house). Four weeks post-surgery, mice were perfused with 4% paraformaldehyde (PFA) and processed for immunohistochemistry experiments.

For the *in vitro* electrophysiological experiments, C57BL/6J mice (p28-35) were injected with AAV2/1-EF1α-DIO-hChR2 (H134R)-mCherry (100 nl) (2.21e12 GC/ml, produced in-house) bilaterally in the PPN together with AAV-Retro-pAM-Cre (6.30e12 GC/ml, produced in-house) (200 nl) bilaterally in the SNr. Vglut2-Cre mice (p28-35) received bilateral PPN injections of 100nl of AAV-EF1α-DIO-hChR2 (H134R)-EYFP (7.9×10e12 GC/ml, Penn Vector Core) or 100 nl of AAV1-EF1α-DIO-hChR2 (H134R)-EYFP together with bilateral VM injections of 150 nl red retro-beads (Lumafluor). Mice were processed for electrophysiological recordings four weeks post-surgery.

For transsynaptic tracing experiments, a mix (200 nl) of anterograde transsynaptic AAV1-hSyn-Cre (4.98e12 GC/ml, produced in-house) and AAV-EF1α-DIO-EYFP were injected unilaterally in the PPN and 200nl of AAV1/2-EFLα -DIO-tdTomato (3.49e12 GC/ml, produced in-house)) was injected unilaterally in the SNr of C57BL/6J mice. Mice were perfused for imaging four weeks post-surgery.

For *in vivo* DREADD-mediated chemogenetic experiments, C57BL/6J mice were utilised. For mice in the clozapine-N-oxide (CNO) group and saline group, AAV-Retro-pAM-Cre (200 nl) was injected bilaterally in the SNr, and 100 nl of Cre-dependent AAV encoding the CNO activated excitatory designer receptor exclusively activated by designer drugs (DREADD), AAV2/1-EF1α-DIO-hM3DGq-mCherry (2.16e13 GC/ml, produced in-house) or inhibitory DREADD, AAV2/1-EF1α-DIO-hM4DGi-mCherry (2016e13 GC/ml produced in-house) was delivered bilaterally into the PPN. For mice in the CNO control group, AAV-retro-pAM-Cre (200 nl) was injected bilaterally in the SNr, and 100 nl of AAV2/1-EF1α-DIO-tdTomato was delivered bilaterally in the PPN. Mice recovered from surgery for three weeks prior to the behavioral training.

For *in vivo* optogenetic experiments, Vglut2-Cre mice (p28-35) received bilateral PPN injections of 100nl of AAV-EF1α-DIO-hChR2 (H134R)-EYFP (Penn Vector Core) (7.9×10e12 GC/ml) together with dual fiber-optic cannulas (Doric lenses) implanted above the SNr. For *in vivo* optogenetic combined with chemogenetic experiments, a triple viral strategy was applied. Vglut2-Cre mice (p28-35) received bilateral PPN injections of 100 nl of AAV1-EF1α-DIO-hChR2 (H134R)-EYFP together with bilateral SNr injections of 200nl AAV2/1-EF1α-DIO-hM4DGi-mCherry and bilateral VM injections of 150 nl AAV-Retro-pAM-Cre. Dual optical fibers were implanted above the SNr after viral injection. Jeweler’s screws and dental cement were used to secure the fibers to the skull.

### PD mouse model generation

6-hydroxydopamine (6-OHDA, Sigma) was prepared immediately prior to surgeries. The sterile DPBS solution containing ascorbic acid (0.2%) was used as the vehicle to dissolve 6-OHDA. In order to inject a final concentration of 12 μg into the right striatum, a working stock of 6 mg/ml was made injecting a final volume of 2 μl.

For *in vivo* optogenetic experiments, C57BL/6J mice (p28-35) were first injected with AAV1/2-EF1α-DIO-ArchT-GFP (100 nl) (1.81e13 GC/ml, produced in-house) bilaterally in the PPN together with AAV-Retro-pAM-Cre (6.30e12 GC/ml, produced in-house) (200 nl) bilaterally in the SNr. 2 weeks post viral injection, mice were anaesthetized using a mixture of ketamine (100 mg/kg, i.p.) and xylazine (10 mg/kg i.p.) and were placed into a stereotactic frame. The lesion was performed using a 5 μl Hamilton syringe to deliver either sterile DPBS-ascorbic acid (0.2%) (Saline) or 6-OHDA (12 μg) at the following coordinates relative to bregma: AP: +0.5 mm; ML: −2.0 mm; DV: −3.0 mm into the right dorsal striatum (CPu) according to the stereotaxic atlas. After drilling a 1 mm burr-hole in the skull, a 2 μl volume of solution was infused at the target site at the rate of 0.2 μl per minute. The needle was held in place for at least 5 minutes after injection to minimize retrograde flow along the needle tract. Dual optical fibers were implanted above the SNr after 6-OHDA/Saline injection. Mice were administered a subcutaneous injection of sterile lactated Ringer’s solution and Carprofen analgesia (5 mg/kg, i.p) to facilitate recovery, and were placed on a heat-pad until complete recovery from anesthesia. After 2 weeks’ recovery, mice were processed for rotarod test.

For *in vitro* electrophysiological experiments, C57BL/6J mice (p28-35) firstly received bilateral PPN injections of AAV-EF1α-DIO-hChR2 (H134R)-EYFP (7.9×10e12 GC/ml, Penn Vector Core) (100 nl) together with bilateral SNr injections of AAV-Retro-pAM-Cre (6.30e12 GC/ml, produced in-house) (200 nl). 2 weeks post viral injection, 6-OHDA/Saline was injected into the right dorsal striatum following the same protocol with above. After 2 weeks’ recovery, mice were processed for rotarod test. Slice patch-clamp recordings were performed in 2 - 3 weeks post 6-OHDA / Saline injection.

### Electrophysiology

Animals were deeply anesthetized with isoflurane, perfused transcardially with oxygenated (95% O2 and 5% CO2) ice-cold cutting solution (containing in mM: 194 sucrose, 30 NaCl, 2.5 KCl, 1 MgCl_2_, 26 NaHCO_3_, 1.2 NaH2PO_4_, and 10 D-glucose) and subsequently decapitated. Brains were rapidly removed and coronal brain slices (300 μm, Leica VT1000S vibratome) were prepared in ice-cold cutting solution. Following dissection, slices were placed in oxygenated artificial cerebrospinal fluid (ACSF in mM: 124 NaCl, 2.5 KCl, 2 CaCl_2_, 1 MgCl_2_, 26 NaHCO_3_, 1.2 NaH_2_PO4, and 10 D-glucose, osmolarity ∼315 mOsm/kg) to recover at 32 °C for 30min, and then the slices were kept at room temperature for at least 30 min before undertaking recordings.

Acute brain slices were transferred to a recording chamber and held in place with a platinum harp. Slices were visualized by an upright microscope (BX50WI, Olympus Optical, Japan) fitted with a CCD camera (Michigan City, IN). Fluorescence was detected by an LED illumination system (pE-2, CoolLED, UK) with YFP/RFP filter sets (Olympus, Japan). Whole-cell patch-clamp recordings were made using a Multiclamp 700B amplifier (Molecular Devices). During recordings, slices were kept perfused by a gravity fed system with oxygenated ACSF which was maintained at 30 ± 2°C by an automatic temperature controller (TC324B, Warner, US). Recording pipettes were pulled to a tip resistance of 4–6 MΩ (Narishige PC-10) from borosilicate glass (GC150F, 1.5 mm, Harvard Apparatus, UK). For all the recordings except I-V relationship recordings, pipette internal solution contained (in mM): KMeSO4 135, NaCl 7, HEPES 10, Mg_2_ATP 2, Na_3_GTP 0.3, biocytin 0.3%, pH 7.3 adjusted with KOH; osmolarity ∼290 mOsm/kg. For the synaptic I-V relationship recordings, the following pipette solution (in mM) was used: CsMeSO4 135, NaCl 8, HEPES 10, Mg_2_ATP 2, Na_3_GTP 0.3, Phosphocreatine 7, biocytin 0.3%, pH 7.3 adjusted with CsOH; osmolarity ∼290 mOsm/kg. Electrode offset potentials were corrected prior to giga-ohm seal formation. Liquid junction potentials were not compensated for. The series resistance was typically 10–30 MΩ and was monitored during the experiments. ChR2-expressing axon terminals were light-activated with 5 or 100 ms whole-field LED illumination using blue excitation wavelengths (470 nm) at a constant power of 1.5 mW (0.09 mW for subthreshold stimulation). For voltage-clamp average responses, neurons were held at −60 mV, and at least 10 photostimuli (duration, 5 ms; wavelength, 470 nm) were given at 10 sec intervals. For the I-V relationship study, responses were recorded at holding potentials from −80mV to +40mV with 10 mV increments. For current-clamp recordings, recordings were corrected for bridge-balance and pipette resistance. As the SNr neurons have spontaneous firing even at resting membrane potential, the recorded neuron was injected with a negative current to inhibit the spontaneous activities prior to response recordings in current clamp. Light-evoked response amplitudes were measured from baseline to peak. The latency was defined as the time between the onset of the optogenetic stimulation pulse and the onset of the postsynaptic response (5%). Jitter was the standard deviation of the latencies of each trace from one recording. Rise time was the time from 10% to 90% of the peak of the response. Paired pulse ratio (PPR) was measured as the ratio of the amplitude of the second light-evoked response to that of the first. Data were acquired using AxoGraph X (AxoGraph), filtered at 10 kHz and digitised at 20 kHz using an ITC-18 board (Instrutech, Port Washington, US). Off-line analysis was performed with AxoGraph X.

For drug application experiments, all drugs were bath applied with the following concentrations: tetrodotoxin (TTX, 0.5 μM, Sigma) and 4-aminopyridine (4-AP, 100 μM, Sigma), mecamylamine (10 μM, Sigma), 6-Cyano-7-nitroquinoxaline-2,3-dione (CNQX, 10 μM, Sigma), D-APV (D-(-)-2-amino-5-phosphonvaleric acid) (30 μM, Sigma), Gabazine (10 μM, Tocris), Ifenprodil (5 μM, Sigma), CNO (15μM, AK Scientific).

### Behavioral procedures

All mice were handled for at least 5 days before any behavioral procedures for 5–10 min per day to reduce stress introduced by contact with the experimenter. Mice were placed in the room for behavioral assays for 30 min for room habituation. All chemo- and optogenetic experiments were started 3 - 4 weeks post-surgery. For PD mice, behavioural tests with optogenetic were performed in 2 - 3 weeks post 6-OHDA injection. Following behavioral experiments, mice were sacrificed, brains sectioned, and injection sites were verified using immunochemistry (below). Mice lacking viral expression in target areas were not included in the experimental analysis.

For chemogenetic behavioral experiments, the behavioral experiments were performed 30 min post CNO (1 mg/kg, in 2% DMSO, AK Scientific, CNO group, CNO control) or saline injections (1 ml/kg, in 2% DMSO, saline group). For the optogenetic activated behavioral experiments, trains of light pulses (10 ms pulse at 10Hz, 20Hz or 40 Hz for 1s, 473 nm, light intensity 166mW/mm^2^ at each fiber tip) were delivered via the optical fiber (fiber tip diameter 200μm). For behavioral experiments involving both optogenetic stimulation and chemogenetic inhibition, behavioral experiments were performed with light stimulation initially and then mice were returned to their home cages for 30 min for resting. Following the 30 min rest, CNO was intraperitoneally injected. Behavioural experiments were performed again with the same light stimulation protocols 30 min post CNO injections. For the optogenetic inhibition behavioral experiments, 561nm laser were applied for continuous 30s (light intensity 203mW/mm^2^ at each fiber tip). Each session of the optogenetic behavioral experiments was divided into 3 epochs: pre-laser, laser and post-laser.

### Open field test

#### Open field test with chemogenetics

The behavioral experiments were performed 30 min post CNO or saline injections. The open field test (activity monitoring system) was applied to monitor the motor activity of mice. Mice were placed in a square plexiglas box (27 × 27 cm) and motor activity of mice recorded using the activity monitoring system (Med Associates Inc, USA) for 60mins: the first 25 min were considered for acclimatization and the next 35 min for the analysis. The distance travelled and the velocity were analyzed using the activity monitor 7 software (Med Associates Inc, USA).

#### Open field test with optogenetics

Each mouse was placed in a square arena (30 cm × 30 cm). Movements of mice were recorded with a wireless camera from above (20 fps, resolution of 1024 × 840 per camera view, D-link Wireless N Day/Night Cloud Network Camera, DCS-932L) under different light stimulation protocols. Videos were exported and processed for analysis. To analyze the locomotor behaviors of mice in the open field test, we applied a background subtraction technique combined with segmentation (*42*). Prior to recording each mouse in the open field arena, a video of the empty white arena was captured, which was used as the background. By subtraction of the background, images containing mice only were obtained. Next, applying image segmentation, the location of the mouse in the arena was tracked. Based on the location information, we then obtained the locomotor speed and distance moved by the mice in the open field arena. Locomotion was defined when locomotor speed was larger than 3 cm/s for at least 0.5sec and the remainder was characterized as immobile time (*43*).

##### Rotarod test

An accelerating rotarod (Ugo Basile, Italy) was used to assess motor coordination and balance. Mice were trained to run on a rotational cylinder rod, which slowly accelerated from 5 to 40 rotations per minute over 5 min. It was confirmed that the mice were walking forward rather than just holding onto the rod. For mice from chemogenetic studies, mice were tested on the rotarod 30 min after CNO or saline injection. For PD mice, the rotarod test was performed at 14 days after 6-OHDA administration (*44*). Latency to fall off the rod was recorded. Three trials per day were performed and the average was presented.

##### Treadmill gaits and limb kinematics recording

A customized gait analysis system was developed to acquire and analyze the footprints and joint kinematics of mice. The system is equipped with a transparent treadmill belt (40 x 4cm). A dual position high-speed camera setup was utilized on this system with one camera on the bottom and one on the side (120fps, resolution of 1920×1080 per camera view, EZVIZ S6, China).To achieve a better visualization of the footprints, a dual LED light system was utilized with a red panel on the top acting as the background and a green strip light embedded on the side to illuminate the footprints. Based on this special optical design^42,43^, when the paw landed on the transparent belt, the touching area of the paw will be highlighted and exported for following analysis. Prior to any recordings, mice were trained to walk on the treadmill. A baseline treadmill speed of 15cm/s was applied for all gait recordings. Using the bottom view, the mouse gaits were filmed, and by the side view, the hindlimb joint activities were captured, enabling the limb kinematics of mice to be analyzed.

### Machine learning based data analysis

#### Gait analysis

Gait analysis was performed with a newly developed machine learning based animal multi-pose tracking system, the Social LEAP Estimates Animal Poses (SLEAP) (*45*). Here for the footprint recognition, we utilized the Top-down estimation model of SLEAP. We selected the body center point of the mouse as the anchor and applied the EfficientNet B7 network as the skeleton. Firstly, we trained the EfficientNet network to identify points of interest on the mouse body (including the nose tip, body center point, tail-base, fore paws and hind paws). We selected a set of 837 frames from 16 videos of seven different animals and trained the EfficientNet network with default parameters for 2000 iterations. The model error of the final trained model that we used for footprint analysis was 3.62 pixels with an object keypoint similarity (OKS) score of 0.90823. Based on the footprint information obtained by the SLEAP model, we analyzed a series of gait parameters. A step cycle duration is characterized as the time from when the paw initially treads the belt to the beginning of the next tread of the same paw. A complete step cycle is divided into two phases: stance phase (the period when the paw touches the belt surface) and swing phase (the period when the paw lifts off the surface). Stride length is defined as the distance between two consecutive initial contacts of the paw. According to the gait classification methods described in a previous study ^3^, we grouped the mouse gaits into four different patterns based on the footprint information. Briefly, walk was defined as a pattern of limb movement in which three or four paws were simultaneously in stance phase. Trot was characterized by a pattern of gait in which a diagonal pair of paws were in the stance phase simultaneously while the other pair moved forward synchronously. Gallop or bound are identified as when the two hind paws were in stance phase at the same time. A locomotor stop was considered when the stance phase outlasted 400ms ^11^.

#### Limb kinematic analysis

Similar to the gait analysis, limb kinematics was also analyzed using the SLEAP deep learning markerless tracking system with some modifications to the network. For limb analysis, we trained the EfficientNet B7 network to identify nose tip, body center point, tail-base and the joint points on the hindlimb (iliac crest [IC], hip, knee, ankle, and metatarsophalangeal (MTP]). A dataset of 1052 frames from 22 video-clips of six different animals were selected for training. After 2000 iterations, we obtained the final trained model with a model error of 2.19 pixels and an OKS score of 0.93002. The identified joints were used to quantify the joint angles. The angular variations per step cycle (angle amplitude) were calculated based on the step cycle duration obtained from the bottom view and used to assess the joint function.

### Immunohistochemistry

For neural reconstruction, biocytin (0.3%) was included in the internal solution for electrophysiological recordings. After recordings, brain slices were fixed with 4% paraformaldehyde (PFA) in 0.1M phosphate buffered saline (PBS) overnight at 4 °C. Slices were then washed with 0.1 M PBS (3 × 15 min) and placed in a blocking solution containing 3% bovine serum albumin (BSA), 0.1% saponin, 3% triton X-100 and 0.1M PBS for 2 hours at RT. After blocking, slices were incubated with Alexa Fluor - conjugated streptavidin (1:1000 in blocking solution, Invitrogen) overnight at RT. After a triple wash with PBS, the brain slices were stained with the nuclei marker DAPI (1:500) and then mounted on a glass slide with DABCO, processed for fluorescence visualisation.

For tracing studies, mice were deeply anesthetized with lethabarb (0.1ml/10g) four weeks post-surgery and transcardially perfused with 4% PFA (in 0.1M PBS). Brains were taken and fixed in 4% PFA at RT overnight and 50 μm slices were cut with a vibratome (Leica VT 1000 S). For brain sections from the anterograde and trans-synaptic tracing studies, slices were stained with the DAPI and processed for imaging. For slices from mice of the retro-tracing studies, slices were first incubated with blocking solution for 2 hr at RT, and then incubated with primary antibody anti-Acetyltransferase (1:3000, AB144P, Millipore) for 72 hr at RT. After rinsing, slices were incubated in the secondary antibody - donkey anti-goat Alexa Fluor 647 (1:1000, Invitrogen) overnight at RT. Slices were washed three times with 0.1M PBS and nuclei were stained with DAPI. Slices containing the entire PPN region from rostral to caudal, were selected for imaging. Four PPN images were chosen along the rostro-caudal axis from each mouse according to the Allen atlas (Available from mouse.brain-map.org) and cellular quantification and overlap analysis was conducted. TdTomato positive, Gad67-EGFP positive and ChAT positive neurons in the PPN region were quantified with Image J software.

For mice from PD studies, mice were transcardially perfused with 4% PFA (in 0.1M PBS) and post-fixed overnight. 50 μm slices were cut with a vibratome (Leica VT 1000 S). Slices containing the entire CPu and SNc were processed for tyrosine hydroxylase (TH) immunostaining. Briefly, sections were incubated in a blocking buffer for 1 hour and stained for TH (MAB318 1:1000) overnight at room temperature. After washing, sections were then incubated with Alexa-fluor 555 secondary antibody (Invitrogen). Sections were then washed and counterstained with DAPI, processed for fluorescence visualization.

All sections were imaged with an upright fluorescent microscope (×5, x10 and ×20, Zeiss, Axio Imager) or a SpinSR10 Spinning Disk Confocal (x10, x20, and x40, Olympus) and Z-stack were obtained.

### Data analysis and statistics

GraphPad Prism version 9.0 and Axograph were used for statistical analysis. Group data was expressed as mean ± standard deviation of the mean (SD) unless indicated. Parametric tests were used to test if values passed a normality test, otherwise a non-parametric test was used. For *ex vivo* electrophysiology, paired t-tests were used to compare the responses before and after drug application. For behavioral experiments, mice were randomly assigned to the CNO group and saline group. Behavioural data were analyzed using a one-way ANOVA test with repeated measures followed by a Tukey’s multiple comparisons test (parametric test) or a Dunn’s multiple-comparisons test (nonparametric test) to identify significant groups, as indicated in the figure legend.

## Acknowledgements

This work was supported by grants from the National Health and Medical Research Council of Australia (2009957 to TMW and 2019271 to PS).

