## Supplementary material for "Modulation of pedunculopontine input to the basal ganglia relieves motor symptoms in Parkinsonian mice": Supplental Figures

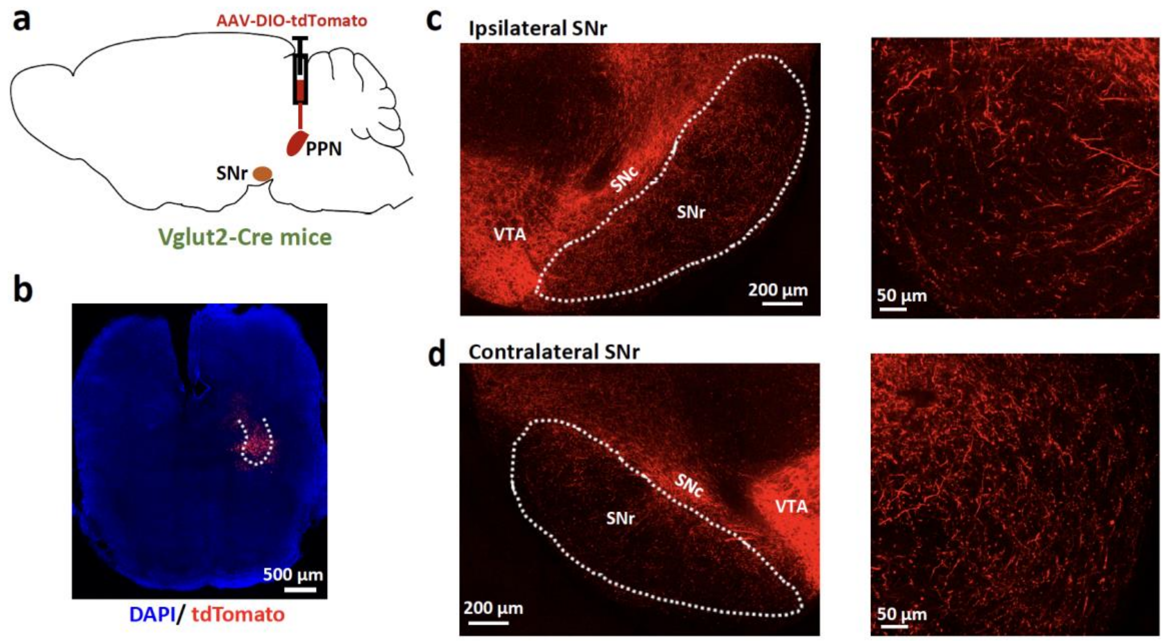


**Figure S1. Glutamatergic neurons in the PPN innervate the SNr.**
**a,** Schematic for labelling glutamatergic neurons in the PPN using AAV- DIO-tdTomato in Vglut2-Cre mice. **b,** tdTomato positive neurons in the PPN (4 weeks post viral injection). PPN border depicted with white dashed line. **c, d,** Glutamatergic axon terminals (red: tdTomato positive; blue, DAPI) in the ipsilateral and contralateral SNr. SNr border depicted with white dashed line. Right panels of **c** and **d** show high magnification of red positive synaptic axons in the SNr. VTA: ventral tegmental area.

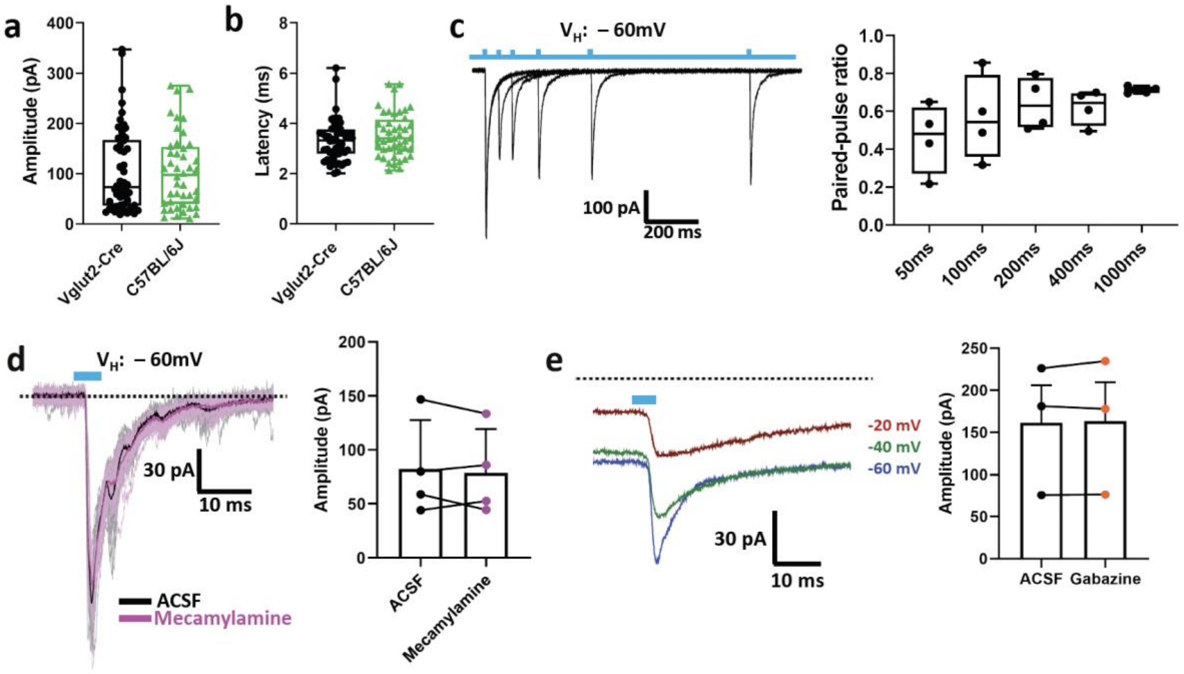


**Figure S2. PPN input to the SNr is glutamatergic.**

**a, b**, Boxplots of the amplitude (**a**) and latency (**b**) of the photo stimulation responses recorded from anterograde labeling (Vglut2-Cre mice, n = 51, error bars represent SD) and retro-labeling studies (C57BL/6J mice, n = 44, error bars represent SD). **c**, Paired-pulse stimulus with intervals ranging from 50 ms to 1000 ms produced depressing synaptic responses in SNr neurons. Right plots show the summary of paired-pulse ratio (PPR) responses (n = 4, error bars represent SD). **d,** Representative traces (left) at -60mV from a SNr neuron in a C57 BL/6J mouse before (black: average of response; light grey: raw traces) and after bath application of 10 µM mecamylamine (purple: average of response; light purple: raw traces) (dashed black line indicates the baseline). Right plots show the summary of amplitudes of responses before and after mecamylamine application (n = 4, error bars represent SEM). **e,** Averaged representative traces showing that light stimulation did not evoke any outward current in SNr neurons of C57 BL/6J mice in voltage clamp holding potential from -60 mV to −20 mV (dashed black line indicates the baseline). Right boxplots show the summary of response amplitudes before and after gabazine application (n = 3, error bars represent SEM).

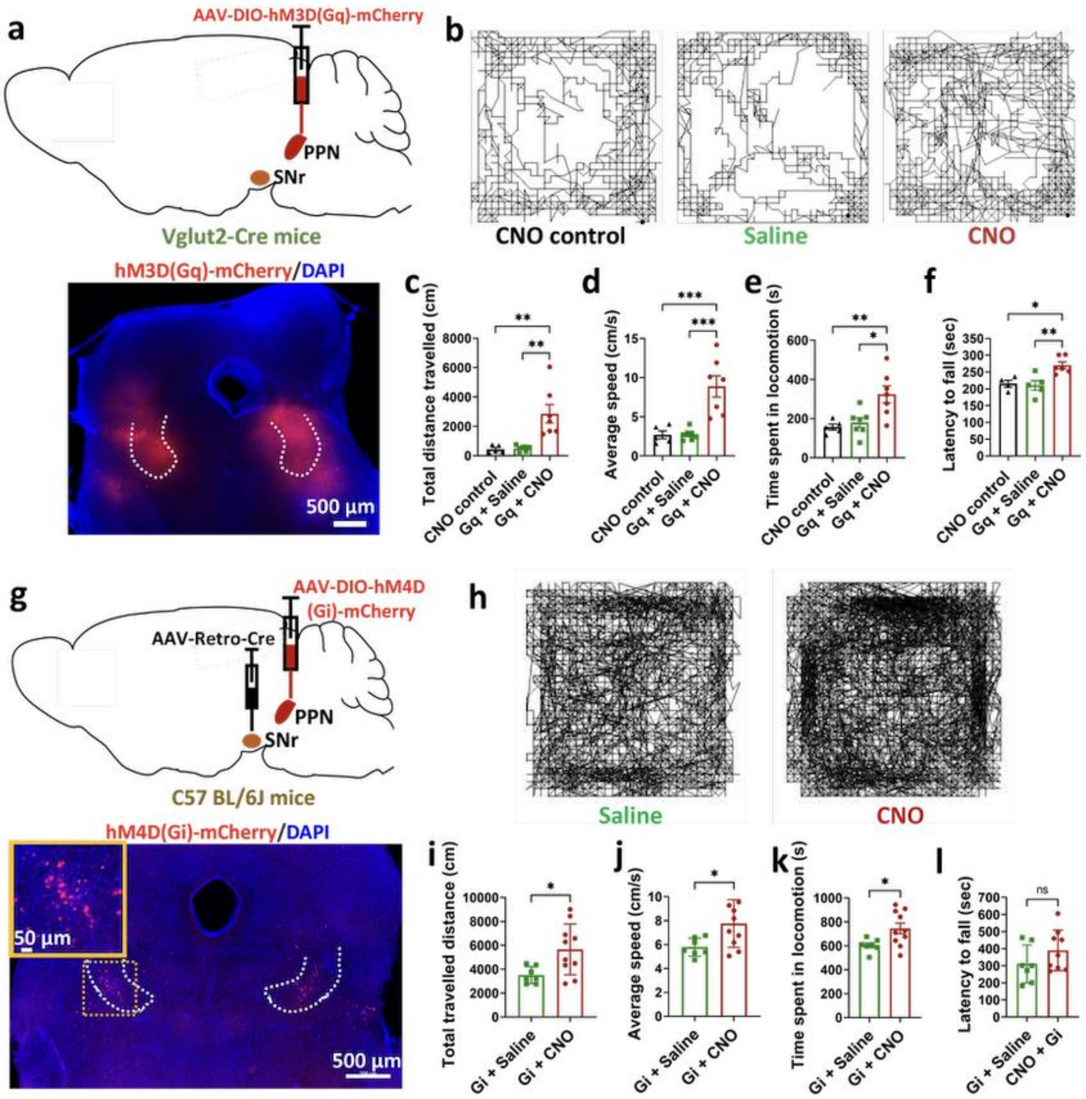


**Figure S3. Increased motor activity caused by chemogenetic activation of non-selective glutamatergic PPN neurons or chemogenetic inhibition of the PPN-SNr projection.**

**a,** Schematic for labelling glutamatergic PPN neurons using AAV-DIO-hM3D (Gq)-mCherry injection bilaterally in the PPN of Vglut2 - Cre mice. Bottom panel shows the hM3D DREADD expressing neurons in bilateral PPN. PPN border depicted with white dashed line. **b,** Representative moving path of mice from the CNO control group (left), Saline group (middle) and CNO group (right) for 60 mins in the open field test. **c-e,** Bar graphs showing quantification of the distance travelled (**c**), average speed (**d**) and time spent in locomotion (**f**) in the open field test of mice in the CNO control group (black, n = 5), saline group (green, n = 5) and CNO group (red, n = 6). **f,** Bar graph showing the latency to fall in the rotarod test of mice in three groups including CNO control group (black, n = 4), saline group (green, n = 5) and CNO group (red, n = 6). **g,** Schematic for retrogradely labelling PPN to SNr projection neurons with hM4D DREADDs using AAV-Retro-Cre and AAV-DIO-hM4D (Gi)-mCherry in C57 BL/6J mice. Bottom panel shows hM4D (Gi) DREADD expressing neurons (red) in bilateral

PPN. PPN border depicted with white dashed line; the yellow boxed insert shows enlargements of the yellow dashed boxed areas (blue: DAPI). **h,** Representative moving path of mice from the saline group (left) and CNO group (right) for 60 mins in the open field test. **i-k,** Bar graphs showing quantification of the distance travelled (**i**), average speed (**j**) and time spent in locomotion (**k**) in the open field test of mice in the saline group (green, n = 7) and CNO group (red, n = 10). Error bars represent SE. **l,** Bar graph showing the latency to fall in the rotarod test of mice in three groups including the saline group (green, n = 7) and CNO group (red, n = 9). For all panels, error bars represent SEM. * $p < 0.05$, ** $p < 0.01$, *** $p < 0.005$. **** $p < 0.001$. “ns” indicates no significant difference.

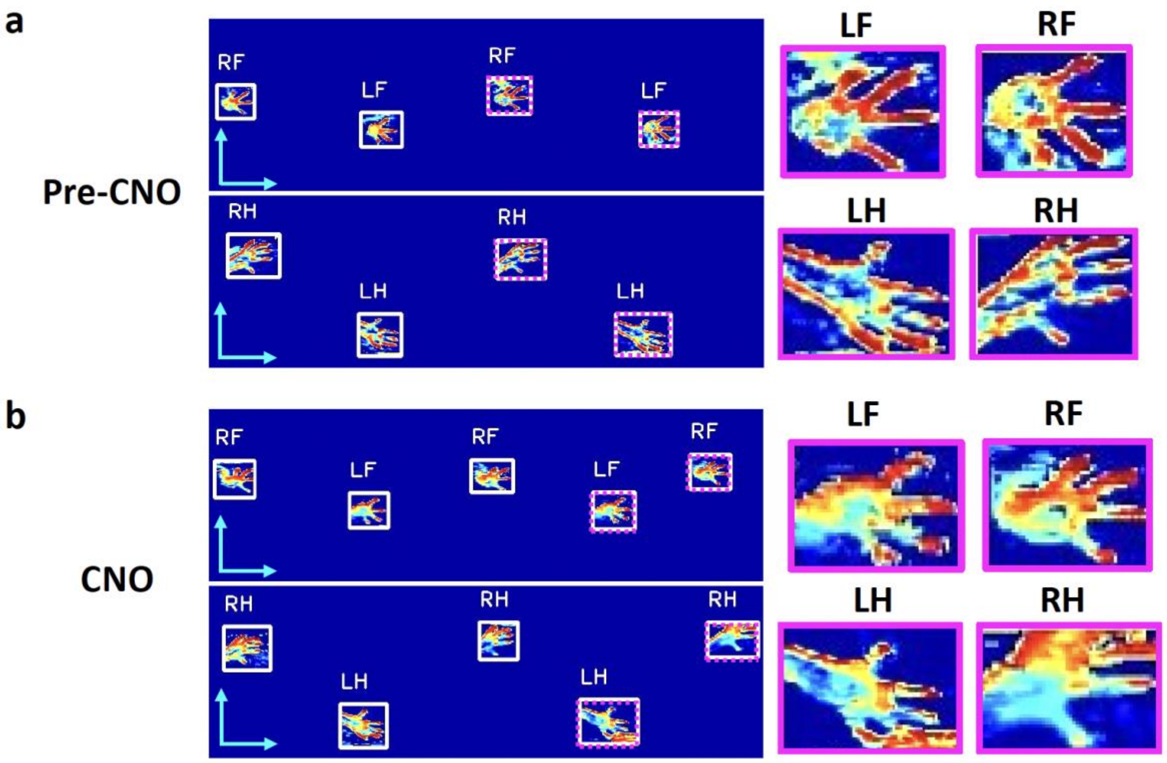


**Figure S4. Representative footprints from a mouse in the CNO group; activation of the PPN input to the SNr increases the footprint.**
**a,** Representative footprints before CNO application. **b**, Representative footprints after CNO application. Scale bar, 1 cm. Purple boxed enlargements are high magnification images of each paw in the dashed boxes.

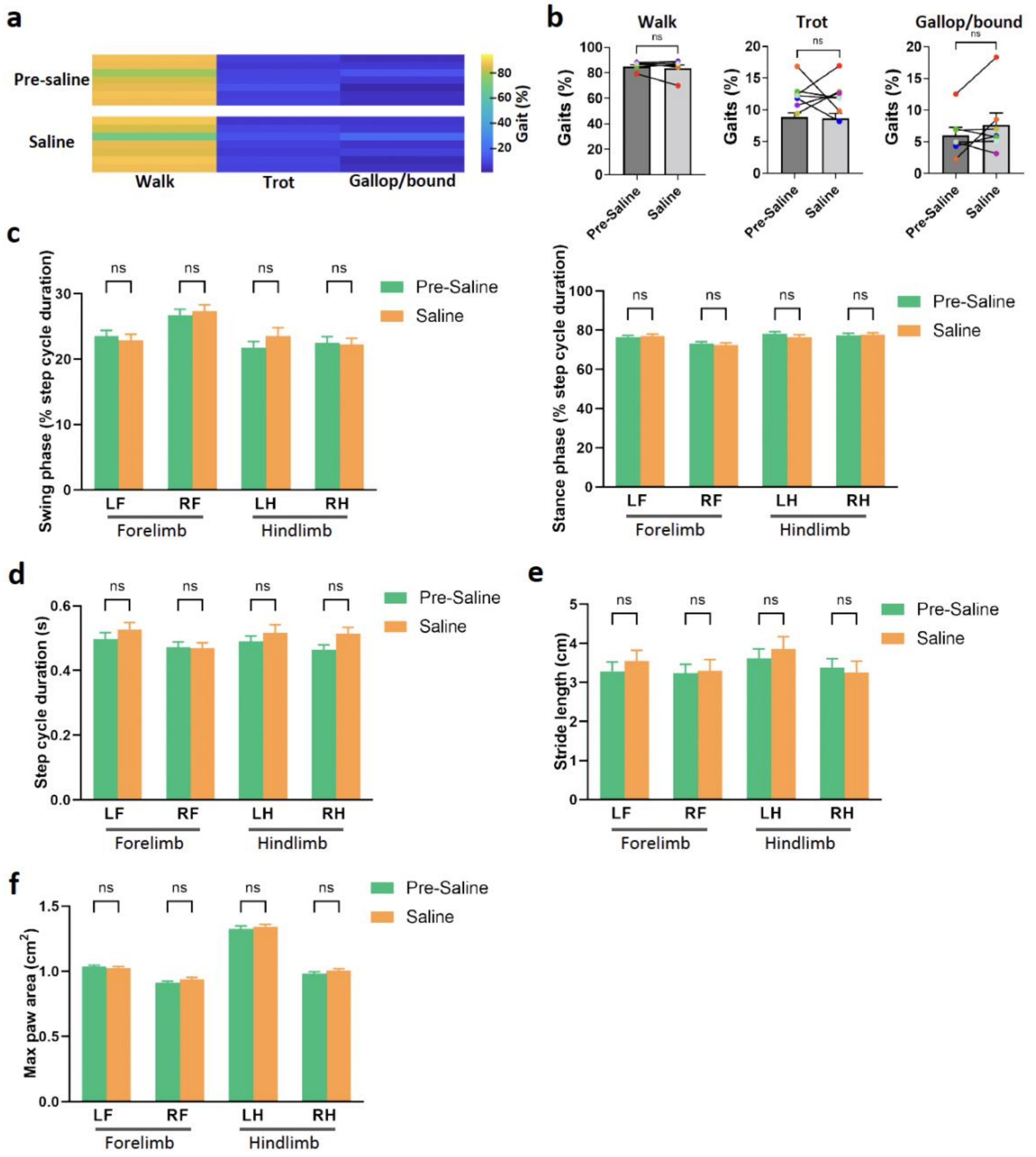


**Figure S5. Control data showing no effect of saline in the excitatory DREADD group.**
**a,** Locomotor gait patterns pre and post CNO injection with individual mice in separate horizontal lines (n = 7). **b**, Bar graphs showing quantification of different gaits (walk, trot and gallop/bound) before and after saline application (n = 7). **c-f**, Mice in the Saline group showed no changes in gait dynamics, including swing phase and stance phase (**c**), step cycle duration (**d**), stride length (**e**) and max paw areas (**f**). For all panels, error bars represent SEM. “ns” indicates no significant difference.

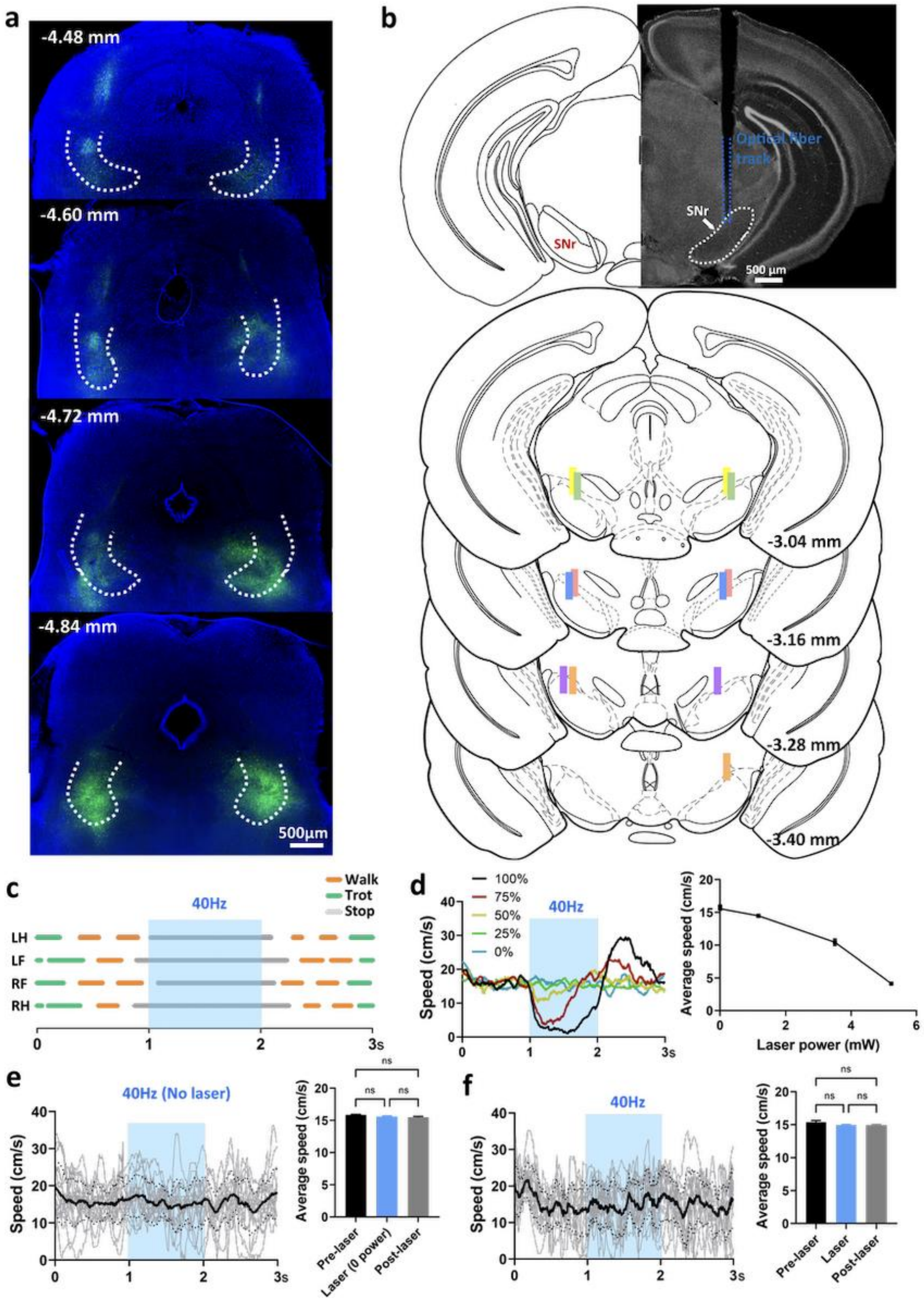
a
-4.48 mm
-4.60 mm
-4.72 mm
-4.84 mm
500μm
b
Optical fiber track
SNr
SNr
500 μm
-3.04 mm
-3.16 mm
-3.28 mm
-3.40 mm
c
Walk
Trot
Stop
40Hz
LH
LF
RF
RH
0
1
2
3s
d
100%
75%
50%
25%
0%
40Hz
Speed (cm/s)
Average speed (cm/s)
Laser power (mW)
e
40Hz (No laser)
Speed (cm/s)
Average speed (cm/s)
ns
Pre-laser
Laser (0 power)
Post-laser
f
40Hz
Speed (cm/s)
Average speed (cm/s)
ns
Pre-laser
Laser
Post-laser

**Figure S6. Verification of viral expression in the PPN and placements of optical fibers in the SNr in Vglut2-Cre mice.** **a,** Representative fluorescence images showing the ChR2 expression in the PPN from rostral to caudal brain sections. **b,** Optical fiber tracks in the SNr. Top panel shows a representative image of the SNr with optical fiber track. Bottom panels depict the location of the optical fiber tip in the SNr along the rostro-caudal axis (summarized from 6 mice, each color represents a single mouse). **c,** Representative gait diagrams from a mouse and the impact of 40 Hz laser stimulation. Shown are gait patterns during three episodes (Pre-laser, Laser and Post laser). Filled bar represents the stance phase and gap represents the swing phase. Blue region depicts the laser stimulation period. **d,** Real time speed during three episodes (Pre-laser, Laser and Post laser) upon 40 Hz laser stimulation with a range of laser powers (n = 5). Right panel show the average speed during three episodes upon different laser powers of 40 Hz stimulation (n = 5, error bars represent SEM). **e**, Locomotor speed upon 40 Hz (no laser) optogenetic stimulation (n = 5, error bars represent SEM). Left panels depicts the real-time speed (light grey: single trials; black: average) and right panel shows the average speed during three episodes. **f**, Locomotor speed in control mice upon 40 Hz laser stimulation (n = 4). Control mice had YFP expression but not ChR2. Left panels depicts the real-time speed (light grey: single trials; black: average) and the right panel shows the average speed during three episodes.

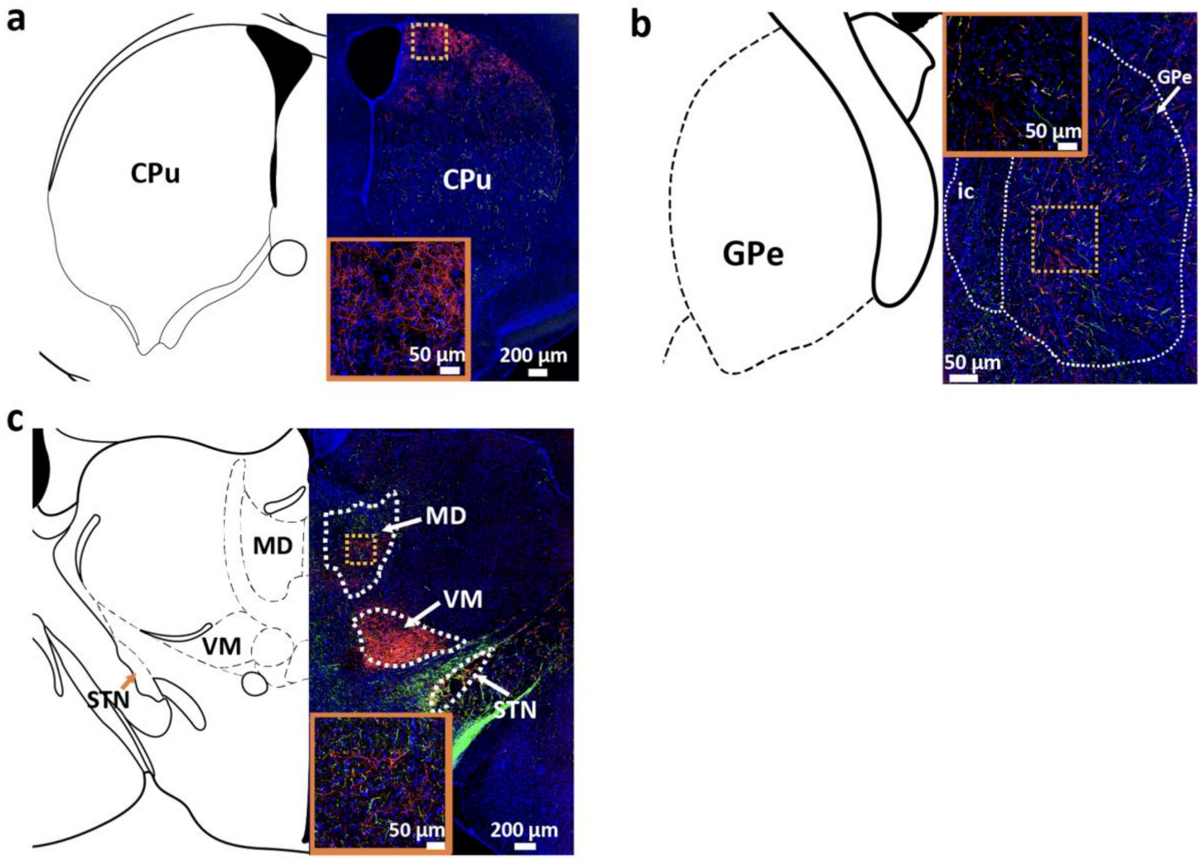


**Figure S7. SNr neurons that receive PPN input project to a number of motor regions..**

**a-c,** Regions that also contain terminals (red) of PPN-innervated SNr neurons include the striatum (corpus striatum, CPu) (**a**), the external globus pallidus (GPe) (**b**) and the medial dorsal thalamus (MD) (**c**). ic, internal capsule; VM: ventral medial thalamus; STN: subthalamic nucleus. Inserts show enlargements of the yellow dashed boxed areas (blue: DAPI).

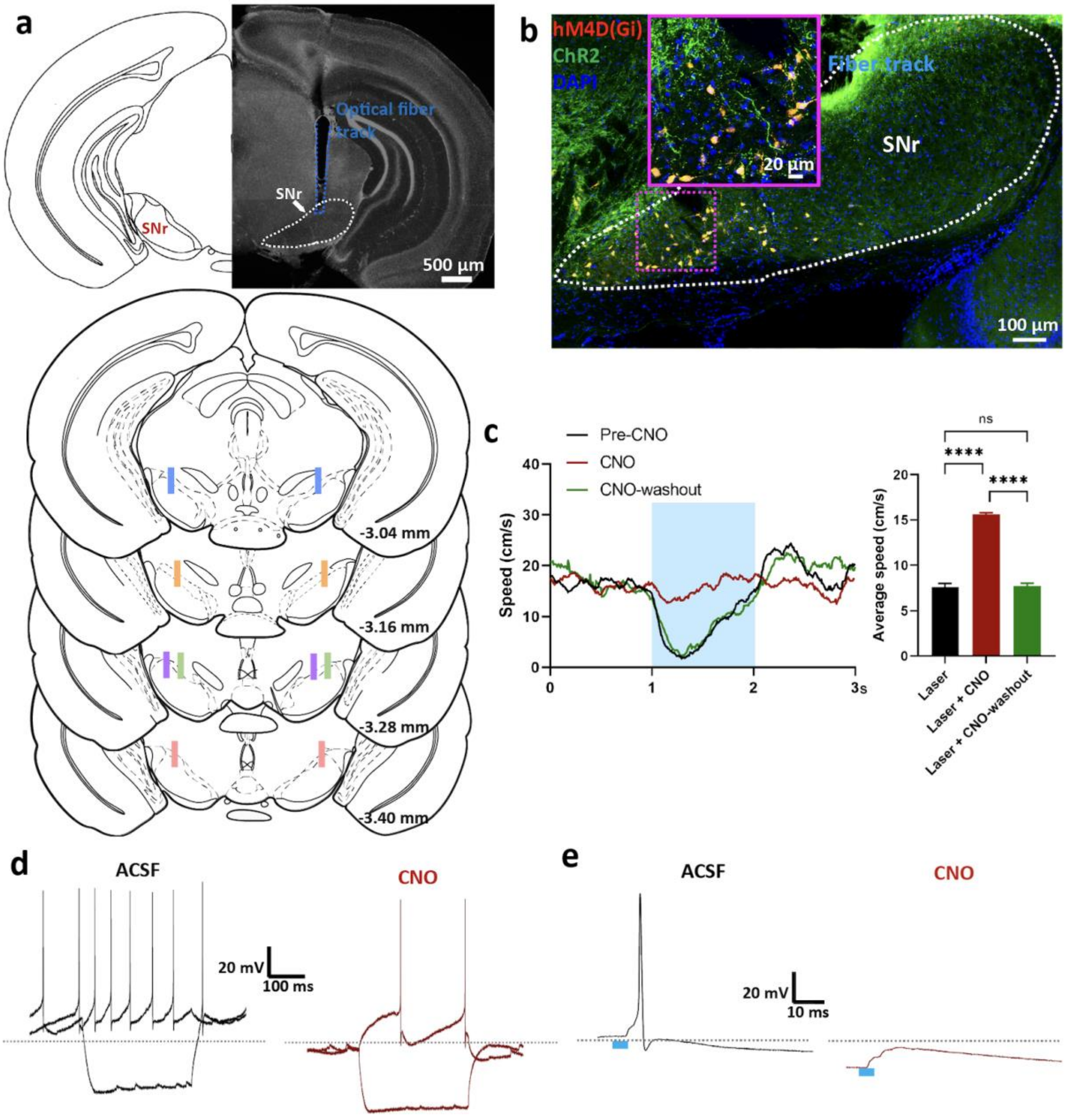


**Figure S8. Histological and electrophysiological verification of hM4DGi expression in SNr neurons, and placement of optical fibers.**

**a,** Optical fiber tracks in the SNr. Top panel shows a representative image of the SNr with the optical fiber track. Bottom panels depict the location of the optical fiber tip in the SNr along the rostro-caudal axis (summarized from 5 mice, each color represents a single mouse). **b,** A representative fluorescent image showing the hM4DGi (red) expression in the SNr from a mouse in the optogenetic combined with chemogenetic behavioral studies. Green: ChR2. Blue: DAPI. **c,** Locomotor speed upon 40 Hz laser stimulation during pre-CNO application, CNO application and CNO wash-out periods. Left panels depict the real-time speed and right panel show the average speed upon laser stimulation in three episodes (n=4, Pre-CNO, CNO application and CNO wash-out). **d,e** Electrophysiological evidence confirming the effect of the CNO. **d**, CNO application decreased the firing rate of the SNr neuron expressing hM4DGi. **e**, CNO application suppressed the AP of the SNr neuron response to light stimulation (blue bar: 5 ms light pulse of 470nm) of the PPN ChR2 terminals.

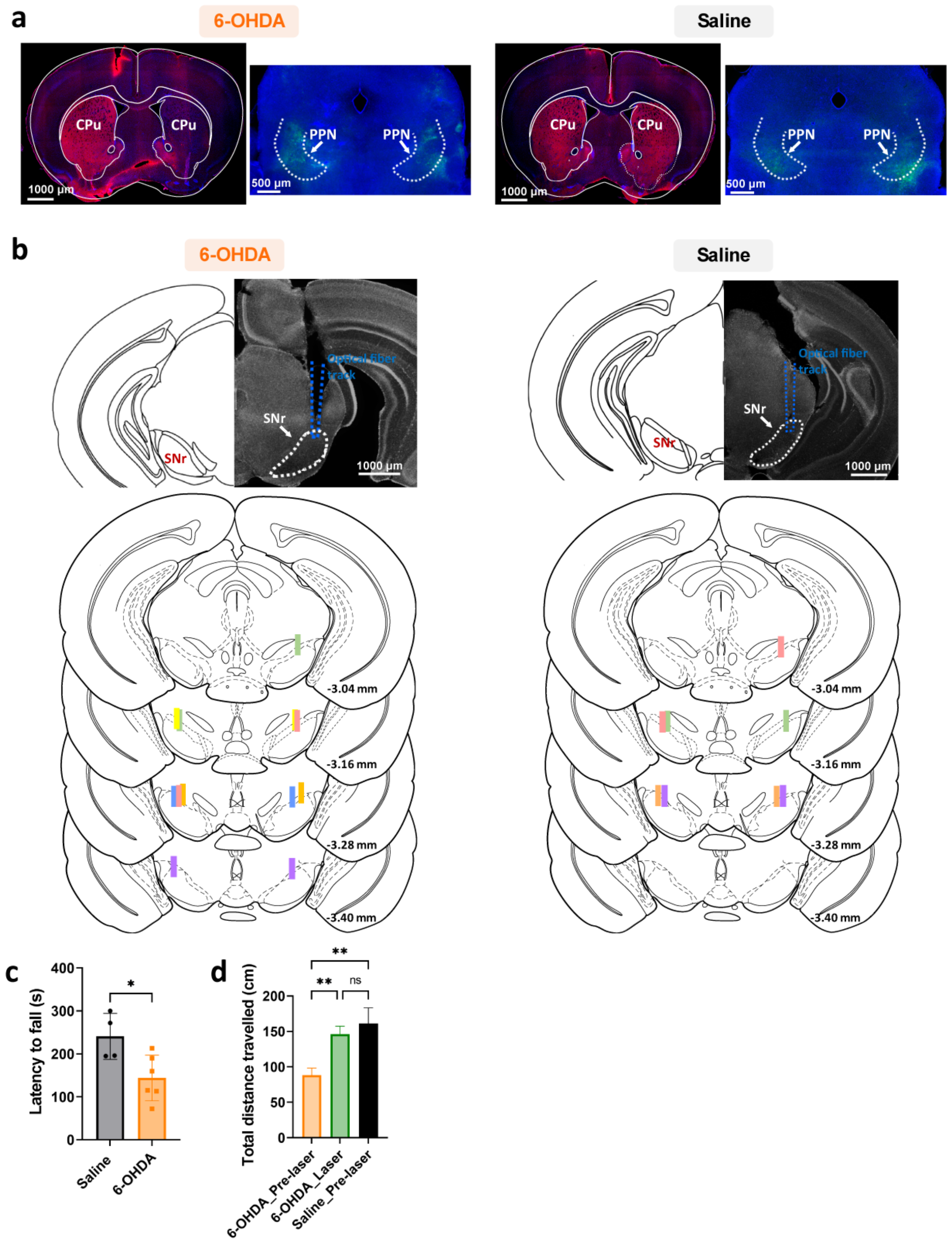
a
6-OHDA
Saline
CPu
CPu
PPN
PPN
1000 µm
500 µm
b
6-OHDA
Saline
SNr
Optical fiber track
1000 µm
-3.04 mm
-3.16 mm
-3.28 mm
-3.40 mm
c
Latency to fall (s)
400
300
200
100
0
Saline
6-OHDA
*
d
Total distance travelled (cm)
200
150
100
50
0
6-OHDA_Pre-laser
6-OHDA_Laser
Saline_Pre-laser
**
**
ns

**Fig. S9 Histologically verified 6-OHDA lesion, viral expression and optical fiber placements of 6-OHDA injected and saline injected mice**

**a,** Fluorescent images showing TH labeled (red) dopaminergic neuron terminals in the CPu and ArchT-YFP expression in the PPN of a representative 6-OHDA-treated animal and saline injected animal, verified the 6-OHDA lesion. Blue: DAPI. **b,** Optical fiber tracks in the SNr of 6-OHDA treated (left, n = 6) and saline treated (right, n = 4) animals. Top panel shows a representative image of the SNr with optical fiber track from a. Bottom panels depict the location of the optical fiber tip in the SNr along the rostro-caudal axis (each color represents a single animal). **c,** Rotarod test data of mice 2 weeks post 6-OHDA (n = 6) or saline (n =4) injection. **d,** Laser stimulation (561nm) reversed the loss of motor activity in PD mice (n =6) to a normal level, the baseline of the saline mice (n = 4).

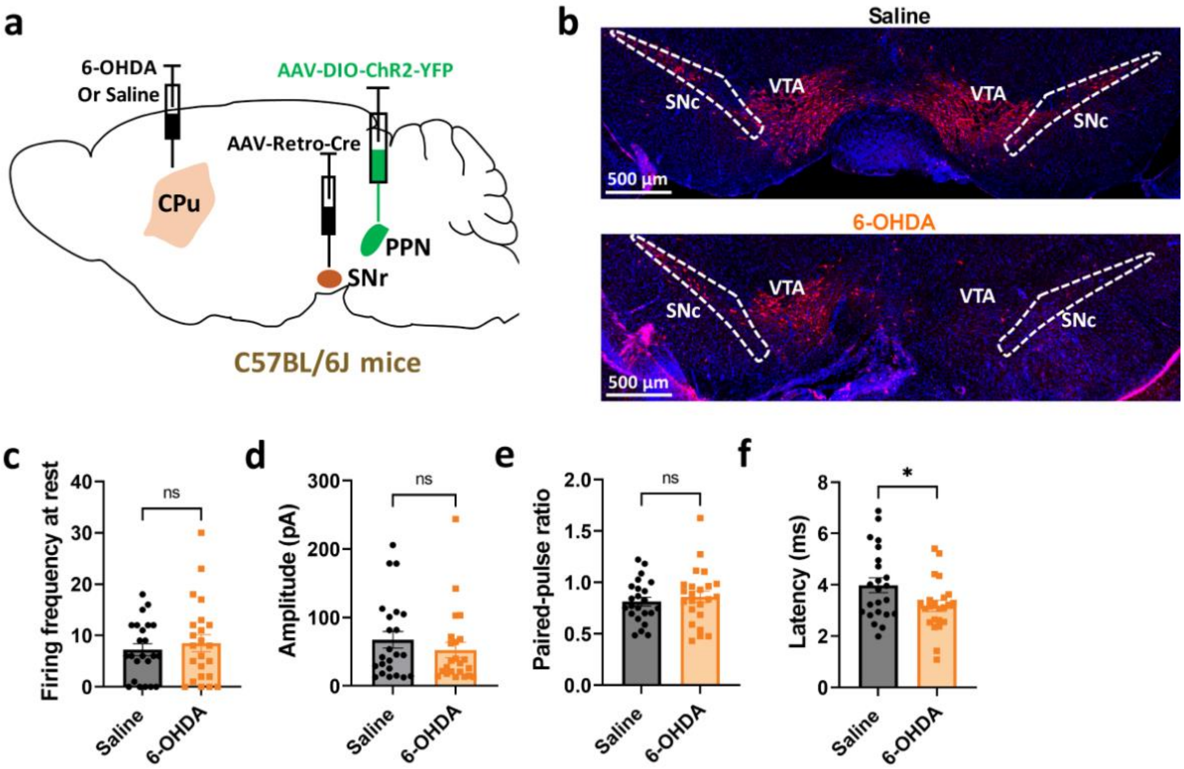


**Fig. S10 Electrophysiological properties of the PPN to SNr projection in PD mice do not change.**

**a,** Schematic for labelling PPN to SNr specific projection neurons with ChR2 in PD mice. AAV-Retro-pAM-Cre was injected bilaterally into the SNr combined with bilateral PPN injection of AAV-DIO-ChR2-mCherry in C57BL/6J mice. 6-OHDA (saline for control mice) was injected in the right CPu to create the PD mouse model. **b,** Fluorescent images show the loss of dopaminergic neurons (labelled with TH staining, red) in right SNc post 6-OHDA injection. Blue: DAPI. **c,** Firing frequency at rest of recorded SNr neurons from 6-OHDA treated (n = 23 cells from 5 mice) and saline treated mice (n = 23 cells from 5 mice). **d - f,** Bar graphs showing the amplitude (**d**), paired pulse ratio (**e**) and latency (**f**) of the photo stimulation evoked responses recorded from SNr neurons of 6-OHDA treated (n = 23 cells from 5 mice) and saline treated mice (n = 23 cells from 5 mice). For all panels, error bars represent SEM. * $p < 0.05$. “ns” indicates no significant difference.
